# EXOSC3 S1-domain variants implicated in PCH1b alter RNA exosome cap subunit abundance and thermal stability disrupting rRNA processing and targeting of AU-rich mRNA

**DOI:** 10.1101/2025.05.31.657176

**Authors:** Avery M. Runnebohm, H.R. Sagara Wijeratne, Monica P. Barron, Whitney R. Smith-Kinnaman, James D. Rooney, Sarah A. Peck Justice, Lauryn A. Cureton, Annalise Holland, Homa Ghalei, Stephane Pelletier, Emma H. Doud, Jonah Z. Vilseck, Amber L. Mosley

## Abstract

Missense variants in EXOSC3, an RNA exosome subunit, have been identified in patients with PCH1b. We investigated three missense variants in the S1 domain of EXOSC3, including one variant of uncertain significance (VUS) and two pathogenic variants (hence S1 variants). EXOSC3 S1 variant cell lines were generated using CRISPR-Cas9 resulting in widespread proteome changes including decreases in some RNA exosome subunits paired with increases in the catalytic subunit DIS3. Thermal stability, analyzed by PISA, revealed extensive destabilization of RNA exosome cap subunits and the cap-associated exonuclease EXOSC10. Functionally, S1 variants altered rRNA processing with corresponding protein compensation observed in rRNA processing proteins outside the RNA exosome. Exogenous overexpression of EXOSC3 rescues many molecular defects caused by S1 variants suggesting that protein destabilization and turnover strongly contribute to molecular defects. Overall, our findings define the mechanisms through which cells respond to EXOSC3 S1 variant disruption of RNA processing homeostasis.

## INTRODUCTION

High-throughput DNA and RNA sequencing have been pivotal in identifying and understanding genetic variants related to human diseases (1–3). Variants observed through clinical genetic testing are classified for pathogenicity based on professional society guidelines and available evidence to guide clinicians in diagnosis and treatment (4). The rates of clinical genetic testing to identify sequence variants are rapidly outpacing functional follow up studies at the protein level, leading to increased accumulation of uncharacterized variants of uncertain significance (VUS). VUS do not have sufficient evidence to support pathogenicity due to factors such as the rarity of the sequence variant, the newness of the associated disease, and/or no established functional link(s) between the variant and the disease, making it difficult for clinicians and patients to understand disease basis and potential for therapeutic intervention.

High-throughput methods for analyzing and comparing functional consequences of VUS and other genetic variants at the protein level could improve our understanding of the proximal molecular changes associated with missense variants in many human diseases.

Several variant alleles of EXOSC3 have been associated with pontocerebellar hypoplasia type 1b (PCH1b) although few have been classified as pathogenic with many remaining as VUS. PCH1b is a rare, neurodegenerative disease characterized by underdevelopment of the pons and cerebellum with motor neuron degeneration in the anterior horn (5,6). PCH1b patients display severe motor impairment and muscle weakness, and most patients do not survive through adolescence (7,8). Genome and exome sequencing of PCH1b patients have identified several variants of the EXOSC3 gene that are associated with PCH1b, some of which are described in Figure S1A (7–12). Clinical evaluation of PCH1b patients with different EXOSC3 variant alleles has shown genotype-phenotype correlations, where certain variants result in more severe symptoms of disease and shorter life expectancies (7). Patients with homozygous G31A variants display the most severe symptoms and do not survive through infancy, while patients with homozygous D132A variants have been observed to experience a range of mild to severe symptoms and in some cases survive into childhood (7,9,13).

Compound heterozygous individuals, typically carrying one D132A variant and another deleterious variant (e.g., frameshift, indel, A139P), usually present with mild to severe symptoms of the disease (5,7,14). The broad range of disease severity caused by EXOSC3 variants raises questions regarding how different variants affect the molecular function of the RNA exosome and cell systems as a whole. Furthermore, PCH1b EXOSC3 variants lead to disease in specific cell types, raising the question of if cellular compensation mechanisms exist in most cell types that can mitigate alterations in EXOSC3 function.

EXOSC3 is a subunit of the RNA exosome, which is a conserved, multi-subunit exonuclease complex that serves as the major 3’-5’ exoribonuclease in the cell required for RNA quality control, processing, and degradation (reviewed in (15–18)). The RNA exosome complex consists of a 3-subunit cap (EXOSC1-3) and a 6-subunit ring structure (EXOSC4-9) that work with catalytic exoribonucleases, DIS3 and/or EXOSC10. EXOSC1-9 serves as a scaffold and RNA binding structure, where the RNA is unwound and threaded through to be degraded by one of the exoribonucleases (19). Additionally, the RNA exosome works with a cohort of other proteins and accessory complexes to identify and target RNAs for processing or degradation.

Nuclear-specific cofactors MPP6 and C1D are required for ribosomal (r)RNA maturation and small nuclear (sn)RNA/small nucleolar (sno)RNA processing and have also been shown to help recruit the MTR4 helicase to the exosome (20–23). The TRAMP, NEXT, and PAXT accessory complexes, all of which also incorporate the MTR4 helicase, promote degradation of aberrant transfer (t)RNAs, messenger (m)RNAs, extended transcripts, non-coding (nc)RNAs, and other small RNAs (15,24–27). In the cytoplasm, AU-rich binding proteins (AUBPs) target RNAs to the RNA exosome as a mechanism for translational control (28,29). Numerous other roles for the RNA exosome have also been described (30). With the requirement of a cohort of proteins cooperating to process and/or degrade RNAs, one aberrant exosome component, such as an EXOSC3 variant, could lead to mechanistic defects within many of the RNA processing pathways or could selectively impact precise activities.

To gain a better understanding of the cellular impacts of EXOSC3 variant alleles, we chose to focus on two pathogenic variants, D132A and A139P, and one VUS, G135R, in the S1 RNA binding domain of EXOSC3 for analysis in human cell models generated using CRISPR-Cas9 (7,9,12). Global proteomics analysis of EXOSC3 S1 variant cells revealed significant decreases in RNA exosome subunit protein abundance with largest effects in a subset of complex members. We show that protein abundance compensation occurs in S1 variant cell lines with significant increases in DIS3 and other proteins associated with tRNA synthesis and rRNA processing. Analysis of protein thermal stability revealed decreased stability of EXOSC3 S1 variants and the cap subunit EXOSC1. The exonuclease EXOSC10 was significantly destabilized in the S1 variants, although EXOSC10 protein abundance was largely unaffected.

These findings suggest possible decreased interactions of EXOSC10 with the RNA exosome complex in S1 variant cells. Functionally, we observed changes in rRNA processing that were not fully resolved by the upregulation of various rRNA processing proteins outside of the RNA exosome. Total RNA-Seq analysis revealed similarities in the molecular phenotypes between the D132A and A139P alleles, but comparison of the G135R VUS with the known pathogenic variants revealed a lower extent of RNA abundance changes. All S1 variants studied here showed significant changes in AU-rich RNAs and cap-associated protein cofactors implicated in AU-rich RNA decay. Exogenous overexpression of WT or S1 variant EXOSC3 for 72 hours can rescue specific molecular defects in S1 variant allele cell lines. Overall, these experiments identify and characterize a continuum of functional changes linked to disease-associated EXOSC3 variants. Further characterization of allele-specific variant modulation of proteome-wide thermal stability and abundance has the potential to further enlighten the range of clinical severities observed in PCH1b patients.

## RESULTS

### PCH1b-associated variants and pathogenic models

Disease associated variants of RNA exosome subunits have mainly been studied in yeast, fly, and mouse models (Figure S1A) (31). Our survey of the literature only identified studies in human cell lines that perturbed RNA exosome subunits through various knockdown approaches (27,32–35). While knockdown reduces the amount of protein, sequence variants can alter protein structure and stability, protein-protein interactions, and/or protein function, therefore effects seen by knockdown of gene or protein expression do not necessarily correspond to effects of genome-encoded missense variants. Furthermore, knockdown is transient or short term since EXOSC3 is a common essential gene in human cell lines, so there is insufficient opportunity for system-wide equilibration of molecular mechanisms to compensate for altered function (36). The analysis of disease related EXOSC3 genetic variant models allow us to identify potential system compensation mechanisms. For EXOSC3 S1 variants specifically, the D132A variant has been modeled using yeast and flies, while G135R and A139P EXOSC3 variants have not been modeled previously (31,37–39). Other variants, specifically G31A and W238R, have been extensively studied in yeast and flies, and in one case, G31A has been studied in induced neural progenitor cells from patient fibroblasts (38,40–42). When prioritizing variants to study, we considered variant allele frequency in PCH1b patients and the general population, position in the protein structure, and pathogenicity. EXOSC3 D132A is the most common allele found in the general population (Figure S1A) and is frequently found in homozygous and compound heterozygous PCH1b patients. D132A, G135R, and A139P are all found in a loop between beta strands of EXOSC3 that interacts with EXOSC5 and EXOSC9 in the protein complex (Figure S1B-E). G135R is a VUS, and A139P has been observed in a compound heterozygous individual with D132A. When exploring allele-specific effects of PCH1b variants, analysis of missense changes within the same domain (S1) and loop of EXOSC3 could reveal if changes within the same structural region lead to similar mechanisms while also evaluating the pathogenicity of the VUS G135R.

### PCH1b-associated EXOSC3 S1 variants lead to wide-spread changes across the proteome

For this work, we have generated HEK293T cell lines with EXOSC3 missense variants using CRISPR-Cas9 gene editing making this the first study of EXOSC3 disease-associated variants in a human model system. Homozygous and heterozygous CRISPR-Cas9 edited cell populations were confirmed by Sanger sequencing (Figure S2). Four biological replicates of homozygous S1 variant cell lines were grown up for multiomics analysis. Global proteomics was performed using tandem mass tag-(TMT-) based protein quantitation using a data-dependent acquisition mass spectrometry approach to determine protein abundance changes in the EXOSC3 S1 variants. Compared to control, 1188 proteins decreased, and 1026 proteins increased in abundance in EXOSC3 D132A/D132A cells (Figure 1A, hence D132A). In EXOSC3 G135R/G135R cells, 781 proteins decreased and 806 proteins increased in abundance (Figure 1B, hence G135R). In EXOSC3 A139P/A139P, 1088 proteins decreased and 1207 proteins increased in abundance (Figure 1C, hence A139P). Differential protein abundance analysis of the two pathogenic variants, D132A and A139P, resulted in similar numbers of protein changes, while the VUS, G135R, had fewer proteins changing in abundance. However, all three S1 variants showed significant protein abundance changes in multiple subunits of the RNA exosome complex (Figure 1A-C, colors as indicated).

**Figure 1:**
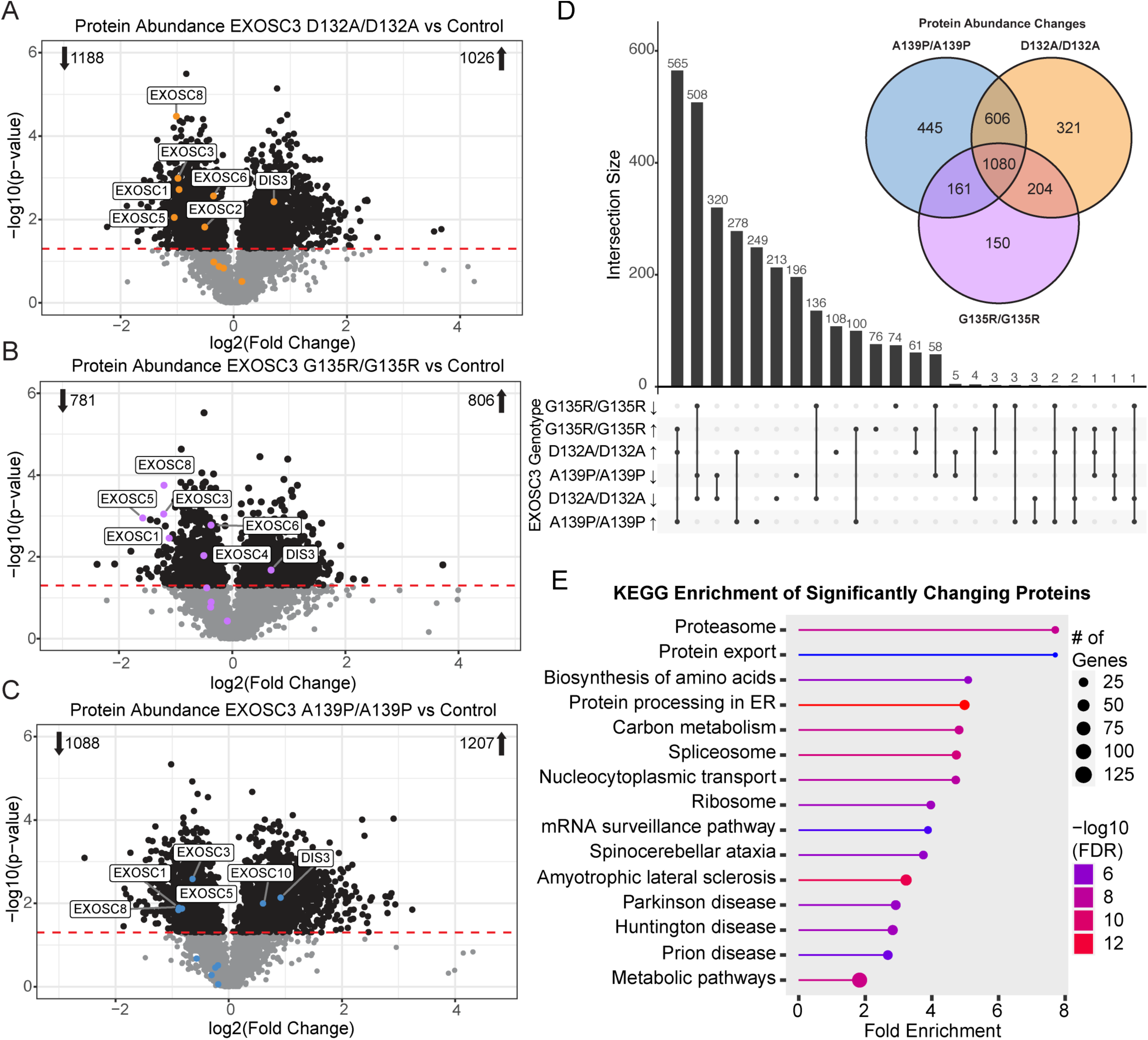
Global proteomics of EXOSC3 S1 variants compared to control. A) EXOSC3 D132A/D132A compared to control. B) EXOSC3 G135R/G135R compared to control. C) EXOSC3 A139P/A139P compared to control. The log2(fold change) for each protein was plotted against the -log10 of the p-value. The red dashed line represents the p-value cut off of 0.05. Each point represents one protein, and black points represent significantly changing proteins. Orange, purple, and blue points represent RNA exosome subunits. Number of proteins significantly increasing or decreasing are shown in the corners of each graph. D) Comparison of protein abundance changes between EXOSC3 variants. UpSet plot and Venn diagram showing overlap of protein abundance changes in EXOSC3 variants compared to control cells. Intersection size is the number of unique or overlapping transcripts. Single dots represent proteins that are unique to that category, and 2 or more dots connected with a line show where there are overlapping protein changes. E) KEGG enrichment analysis of proteins that are significantly changing in abundance in all three S1 variant cells. Pathways and diseases enriched in the 1080 overlapping changing proteins are shown. Marker size represents the number of genes/proteins significantly changed in each pathway/disease. Color represents -log10(FDR). Line length corresponds to fold enrichment.

Overlap analysis indicates high similarity in protein abundance changes between the three EXOSC3 S1 variants. 1080 overlapping proteins are significantly changed in abundance in each of the EXOSC3 S1 variants, with 1073 changing in the same direction (Figure 1D). This UpSet plot shows that there are more protein changes in common between variants than there are changes unique to an individual S1 variants. Gene ontology enrichment analysis of the 1080 proteins overlapping in all three S1 variants shows significant enrichment for changes to the proteasome, spliceosome, and ribosome (Figure 1E). Proteins in the mRNA surveillance pathway are also enriched, which is not surprising since the RNA exosome is directly involved in mRNA quality control, and we show that the RNA exosome subunits as well as some cofactors are impacted by the EXOSC3 variants. Interestingly, several neurodegenerative diseases show statistical enrichment including Parkinson, Huntington, and Amyotrophic lateral sclerosis (ALS).

Most of the exosome cap and core subunits trend towards decreased protein abundance, with significant decreases observed for EXOSC1, EXOSC3, EXOSC5, and EXOSC8 in all three of the EXOSC3 S1 variant cell lines. These subunits also displayed a larger magnitude change in abundance than the other subunits (Figure 1A-C, Figure 2A). EXOSC1, EXOSC3, EXOSC5, and EXOSC8 are all localized to the same face of the protein complex, as shown in the structure determined by Weick et al 2018 (43), suggesting the interaction between these subunits may be altered by these S1 variants. The accessory proteins MPP6 and MTR4 also decreased in cellular abundance, both of which interact with the subset of subunits that are greatly decreasing in abundance. The catalytic subunit DIS3 significantly increased in abundance in each of the EXOSC3 S1 variants, while the catalytic subunit EXOSC10 was only significantly increased in A139P/A139P cells. DIS3 has been proposed to work independently of the complex in other systems and has exosome-independent endonuclease activity in the cytoplasm of human cells (15,44), so this result could suggest compensation for the decrease in the core subunits. We also observe a greater magnitude of decrease of these proteins in G135R/G135R cells compared to the pathogenic variants D132A and A139P. Direct comparison of protein abundance changes between G135R and D132A cells revealed that EXOSC1, EXOSC8, EXOSC10, and MTR4 are significantly lower in cells expressing the VUS G135R (Figure S4). EXOSC3 is decreasing in G135R cells compared to D132A, but this did not reach statistical significance.

**Figure 2:**
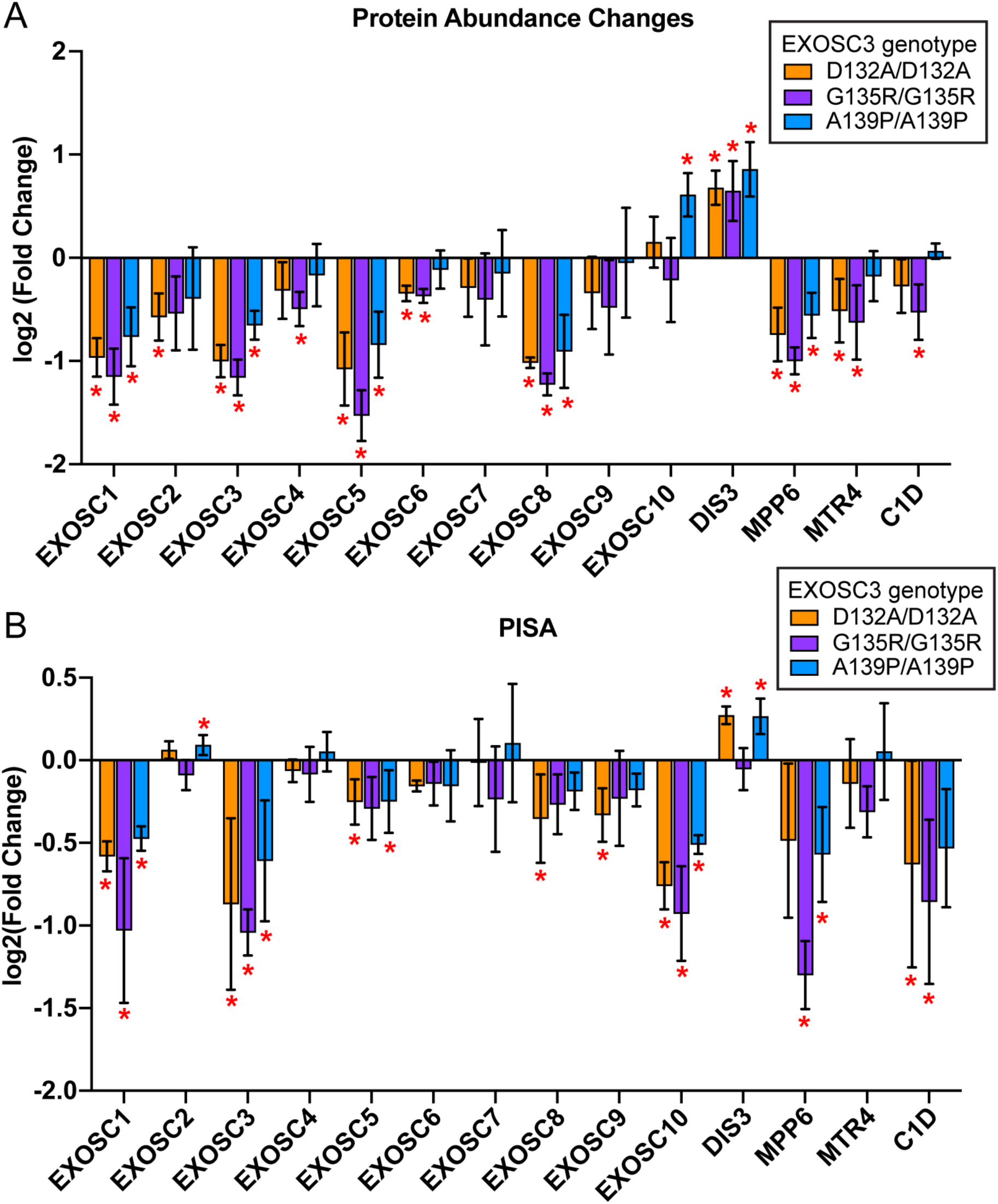
RNA exosome subunit changes in each of the EXOSC3 variants. Fold change in protein abundance (A) and PISA (B) of the RNA exosome subunits and two cofactors, in EXOSC3 variants compared to control. Fold change was calculated by dividing the normalized abundance in the variant samples by the normalized abundance in the control sample for each protein in each replicate (n=4). The log2(fold change) was calculated at the replicate level, and the average log2(fold change) was plotted. Error bars represent standard deviation, and red asterisk represents p ≤ 0.05 (ANOVA, Tukey HSD).

### Thermal stability changes in the RNA exosome indicate allele-specific orphan protein destabilization in the presence of EXOSC3 S1 variants

The EXOSC3 S1 variants may be contributing to RNA exosome dysfunction by altering protein folding, protein-protein interactions, protein modifications, or localization of complex subunits or other proteins important for its function. These biophysical changes to proteins can be detected by analyzing protein thermal stability (45). We used proteome integral solubility assay (PISA) to analyze protein thermal stability (Figure S5C, RNA exosome subunits in color) (45–47). Using PISA, we found that A139P cells had the most proteins significantly changing in stability with 342 destabilized and 543 stabilized (Figure S5C). D132A variant cells resulted in 340 proteins destabilized and 234 proteins stabilized, while G135R cells had 165 destabilized and 81 stabilized (Figure S5A-B). As with protein abundance, we observed lower overall numbers of protein stability changes in the VUS G135R than in the known pathogenic S1 variants. EXOSC1, EXOSC3, EXOSC5, and EXOSC10 were significantly destabilized, although to varying degrees, in each of the EXOSC3 S1 variants (Figure 2B). It is interesting to note that EXOSC10 is destabilized in each of the EXOSC3 S1 variants, but the abundance of EXOSC10 is either unchanged or increasing in S1 variant cells (Figure 2A-B). With several core subunits decreasing in abundance and stability, this could suggest that EXOSC10 protein is present in the cell but is relatively unstable with the steady state changes in RNA exosome complexes.

Conversely, DIS3 was significantly stabilized in both D132A and A139P homozygous variant cell lines (Figure S5C, Figure 2B).

### Differential transcript abundance indicates RNA exosome function is altered by S1 variants

To analyze the functional consequences of EXOSC3 S1 variants on the transcriptome, we performed deep coverage RNA sequencing of total RNA (n=4 per genotype). For each of the S1 variants, we observe a greater number of transcripts decreasing in abundance than increasing relative to control (Figure 3A-C). Changes were broadly observed in both lncRNAs and protein coding mRNAs (Figure S6A, Tables S11-13). Changes in sn/snoRNA, miRNA, and other small non-coding RNAs were also observed (Figure S6A). With 1052 decreasing transcripts and 221 increasing transcripts, D132A/D132A led to more total transcripts changing in abundance than G135R/G135R or A139P/A139P cells (Figure 3A). A greater number of changing transcripts likely suggests a higher degree of RNA exosome dysregulation, indicating that D132A/D132A has a greater impact on RNA exosome function compared to G135R/G135R and A139P/A139P.

**Figure 3:**
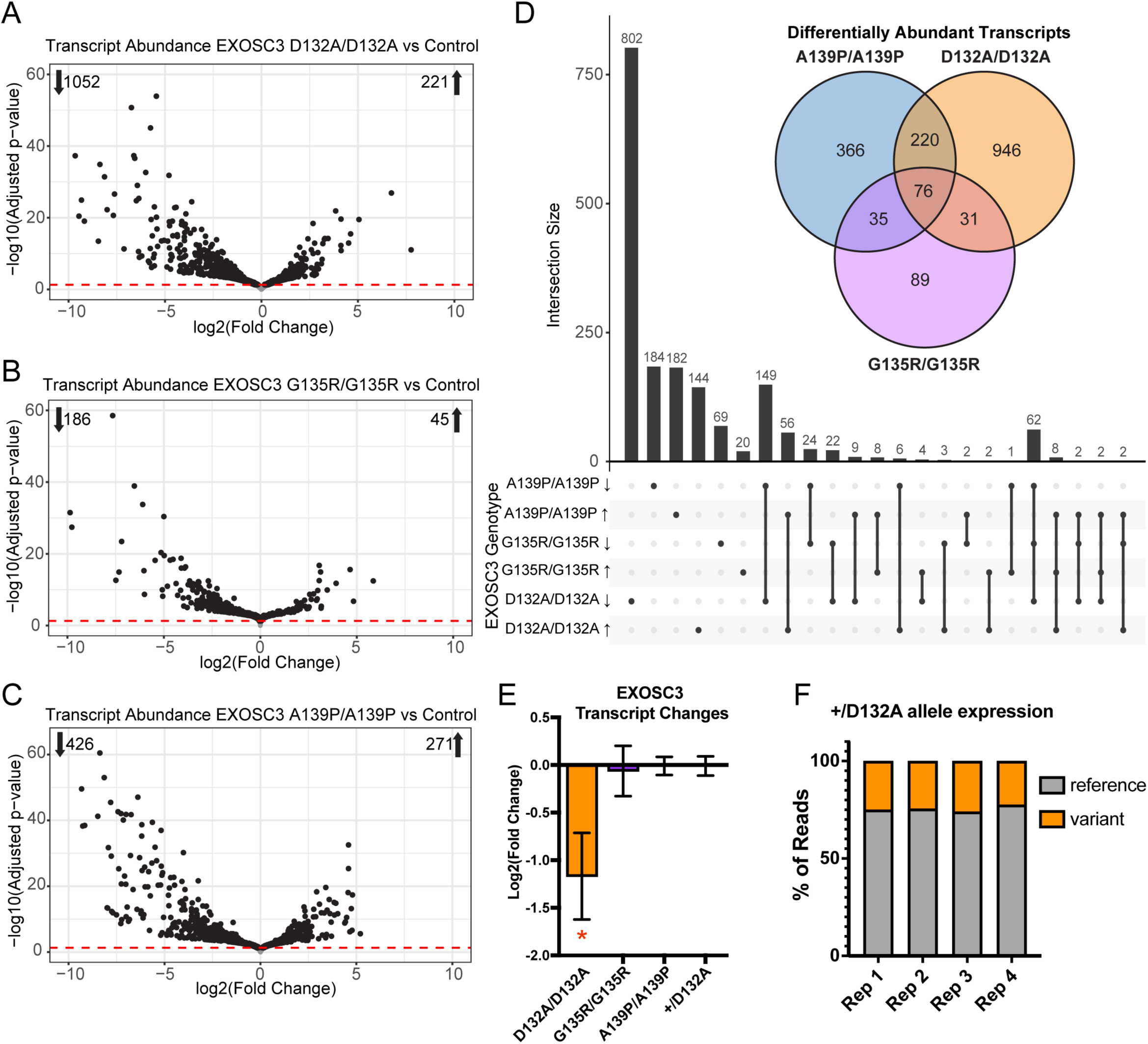
Differentially abundant transcripts in EXOSC3 variant cells. A) EXOSC3 D132A/D132A compared to control. B) EXOSC3 G135R/G135R compared to control. C) EXOSC3 A139P/A139P compared to control. The log2(fold change) for each transcript was plotted against the -log10 of the adjusted p-value. Each point represents one transcript. The red dashed line represents an adjusted p-value cutoff value of 0.05. Number of transcripts significantly increasing or decreasing are shown in the corners of each graph. D) UpSet plot and Venn diagram showing overlap of transcript abundance changes. Intersection size is the number of unique or overlapping transcripts. Single dots represent transcripts that are unique to that category, and 2 or more dots connected with a line show where there are overlapping transcripts. The Venn diagram shows only numbers of changing transcripts while the UpSet plot shows whether the transcripts are increasing or decreasing. E) Transcript abundance changes for EXOSC3. Log2(fold change) for EXOSC3 is plotted for each EXOSC3 variant and the heterozygous D132A variant. Error bars represent standard error; red asterisk = adjusted p ≤0.05. F) Allele specific expression of EXOSC3 in heterozygous D132A cells (+/D132A). Percentage of reads mapping to the reference (gray) or D132A variant (orange) EXOSC3 allele is shown for each of 4 biological replicates.

As EXOSC3 D132A and A139P variants are known pathogenic variants and G135R is a variant of uncertain significance (VUS), we compared these RNA-Seq datasets to identify similarities and differences across the transcriptome. The UpSet plot and Venn diagram comparing the transcripts that are significantly changing indicate that the majority of transcripts changing are unique to genotype, although these S1 variants are all present in the same structural loop (Figure 3D, Figure S1B-E). The greatest overlap in differential transcript abundance is between the pathogenic variants D132A/D132A and A139P/A139P with 149 transcripts decreasing and 56 increasing in both genotypes. There are only 76 total transcripts that are changing across all three of the S1 variants. This contrasts with global proteomics data where there was a high degree of overlap in differentially abundant proteins (Figure 1D). This low degree of overlap could be due to differences in the efficiency of RNA turnover and processing by both the RNA exosome, such as the increased DIS3 protein abundance seen in the S1 variants, and other cellular compensation pathways (Figure 2A). Overall, transcript changes in G135R/G135R cells trend in a similar direction as D132A/D132A and A139P/A139P cells but are not as pronounced in magnitude. Of note, although the S1 variants are not located in regulatory regions, EXOSC3 transcript abundance could be altered by post-transcriptional mechanisms leading to dysfunction. We found that EXOSC3 transcript abundance is decreased in D132A/D132A cells by 50% compared to control, but it is not significantly changing in the other S1 variants (Figure 3E). Transcripts for the other RNA exosome subunits, including DIS3 and cofactors MPP6, MTR4, and SKI, are not changing in any S1 variant datasets (Tables S11-13).

Both homozygous and heterozygous EXOSC3 missense variant cell lines were analyzed by RNA-seq. Cells that are heterozygous have the potential for allele specific expression (ASE) in which there is a bias towards mRNA transcription from one specific allele over the other. ASE is a mechanism influenced by genetic and epigenetic factors that can be used as way to decrease the cellular impact of pathogenic variants (48–52). By analyzing allelic expression in the heterozygous EXOSC3 S1 variant cells, we gain insight into whether this mechanism contributes to the unaffected status of heterozygous individuals. To determine if the heterozygous cell line (+/D132A) displays ASE, deep RNA sequencing was performed (n=4). The number of reads of the EXOSC3 transcripts that contained the reference sequence and variant sequence were compared. In EXOSC3 +/D132A cell lines, we observed 75-78% of transcripts contained the reference sequence, while 22-26% of transcripts contained the variant sequence (Figure 3F). This data shows that EXOSC3 +/D132A cells have an allele specific bias for expression of the reference allele. Although we observe decreased EXOSC3 transcript abundance in homozygous EXOSC3 D132A cells, we do not see an overall change in EXOSC3 transcript abundance in the heterozygous EXOSC3 D132A cells suggesting that the reference allele compensates for the D132A allele (Figure 3E).

### Targeting of AU-rich mRNA is altered in EXOSC3 S1 variant cells

RNA untranslated regions (UTR) are important for RNA binding and transcript regulation. The 5’ UTR is involved in translation regulation, ribosome recruitment and start codon scanning, while the 3’ UTR is important for mRNA stability and localization, regulation by miRNAs, and translation efficiency (53–55). We sought to determine if the UTRs of the differentially abundant transcripts (DATs) were enriched for specific UTR regulatory elements. We used the Atlas of UTR Regulatory Activity (AURA) database for enrichment analysis of our DATs from each genotype (56,57). Several regulatory factors were found to be enriched in DATs for each of the EXOSC3 S1 variants, but alternative polyadenylation (APA) and AU-rich elements (ARE) are present in approximately 40-50% of DATs (Figure 4A, scale to the right). As AREs and AU-rich binding proteins are important for targeting RNAs for degradation by the RNA exosome, enrichment of AREs in the DATs suggests targeting of the RNA exosome is altered by the S1variants. Almost 50% (110 of 231) of the DATs in the VUS G135R/G135R cells have AU-rich regulatory elements, so these DATs were used for enrichment analysis. Using ShinyGo Biological Process, we found that the DATs with AREs are significantly associated with several neural processes including dopamine secretion, neuron differentiation, and neural development (Figure 4B). These data suggest that defects in AU-rich RNA targeting in RNA exosome S1 variants could be a key dysfunction leading to PCH1b.

**Figure 4:**
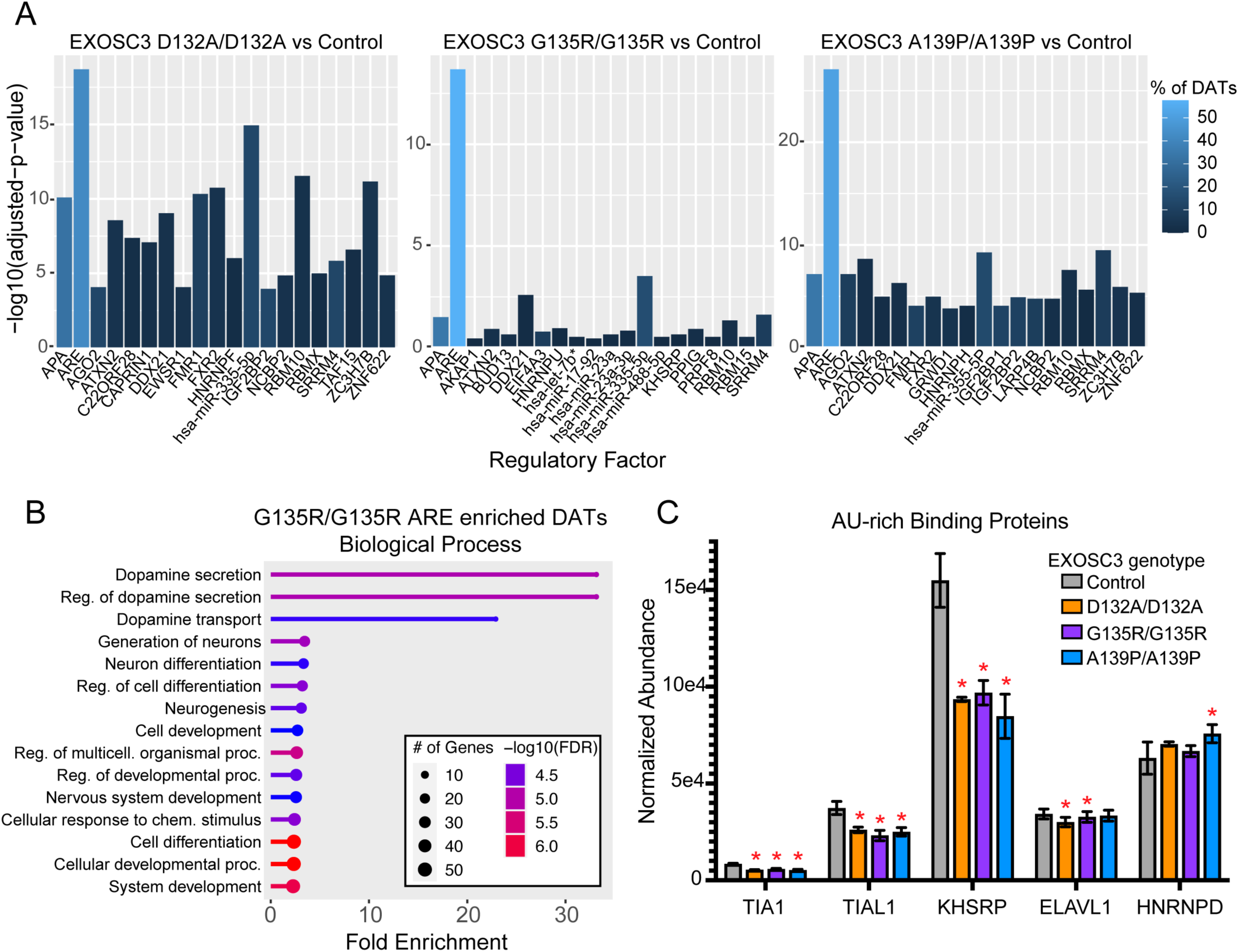
Targeting of AU-rich mRNA is altered by EXOSC3 variants. A) AURA enrichment of UTR regions of differentially abundant transcripts in EXOSC3 variants compared to control cells. UTR regulatory elements enriched in differentially abundant transcripts (DATs) are plotted against -log10 of the adjusted p-value, so increasing size bar has smaller p-value. The color of the bar indicates the percentage of DATs that are regulated by that UTR element. B) Gene ontology enrichment analysis of DATs in the ARE category for EXOSC3 G135R/G135R cells. Size of the point represents the number of genes/proteins significantly changing that are involved in that pathway/disease. Color represents -log10(FDR). Length of line corresponds to fold enrichment. C) Protein abundance changes of AUBPs. Normalized protein abundance of five AUBPs is shown for control and EXOSC3 variant cell lines. Error bars represent standard deviation, and red asterisks represent p ≤ 0.05 (n=4, ANOVA, Tukey HSD).

MPP6 has high affinity for AU-rich RNAs in yeast suggesting a role for MPP6 in AU-rich RNA regulation (23). The dysregulation of AU-rich RNAs is consistent with the decreased abundance and thermal stability of MPP6 observed to some extent in all EXOSC3 S1 variants (Figure 2). AREs are also regulated by non-exosome associated AU-rich binding proteins (AUBPs); therefore, we further assessed protein abundance of AUBPs in the S1 variants. The average normalized protein abundances for significantly changed AUBPs: TIA1, TIAL1, KHSRP, ELAVL1, and HNRNPD are shown for control and S1 variant cells (Figure 4C). We observe statistically significant decreases in abundance for TIA1, TIAL1, and KHSRP in all three of the homozygous EXOSC3 S1 variants compared to control. ELAVL1 is decreasing in EXOSC3 D132A/D132A and EXOSC3 G135R/G135R but is not significantly changing in EXOSC3 A139P/A139P. Conversely, HNRNPD is significantly increased in A139P/A139P cells, and although HNRNPD trends towards increasing in D132A/D132A and G135R/G135R cells it is not significantly changing (Figure 4C). These data further support our transcriptome findings showing that AU-rich RNA targeting is significantly altered in the EXOSC3 S1 variants.

### Ribosomal RNA processing altered by pathogenic variants

Since ribosomal RNA (rRNA) processing is an important function of the RNA exosome, and the protein abundance changes were enriched for “ribosome” (Figure 1E), we analyzed rRNA processing by Northern blot (Figure S7A, n=4). RNA was extracted from control and S1 variant cells using Trizol, and RNA integrity was assessed prior to blotting. For Northern blot analysis and quantification, 7SL was used for normalization, as it is not a target of the RNA exosome. The mature rRNAs 28S, 18S, 5S, and 5.8S show a decreasing trend in EXOSC3 pathogenic variants D132A and A139P, which is significant across replicates for 5.8S rRNA whereas no significant changes were observed for G135R cells (Figure 5A). No significant changes in precursor rRNA abundance were observed; however, some precursors trended down (Figure S7B). With altered rRNA processing in some S1 variants and enrichments in “ribosome” related protein abundance changes across all three S1 variants (Figure 1E), we further analyzed protein abundances of previously defined rRNA processing proteins. Figure 5B shows the abundances for several rRNA processing proteins in control and S1 variant cells.

**Figure 5:**
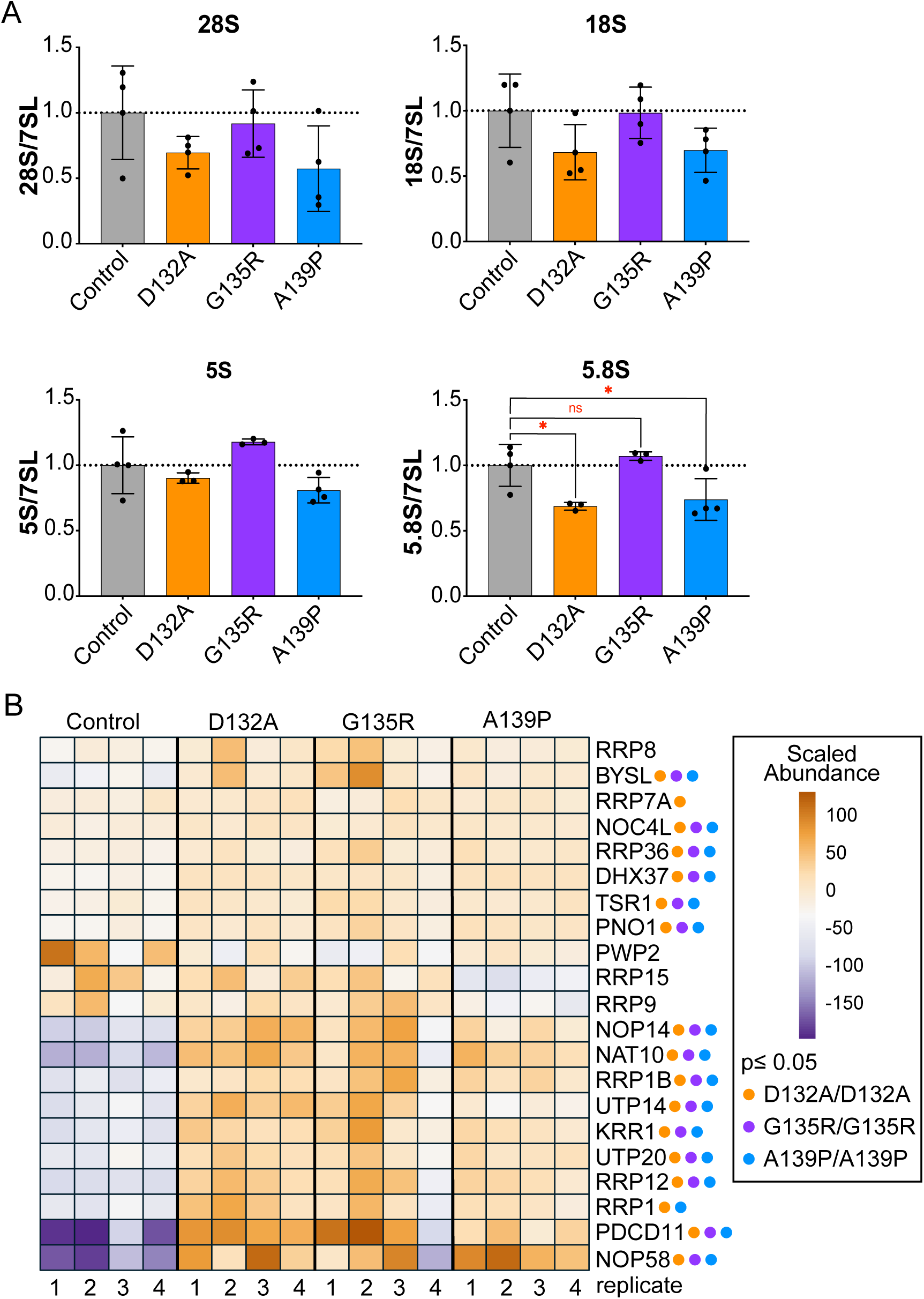
Ribosomal RNA processing is altered in EXOSC3 variant cells. A) Quantification of mature rRNAs from 3-4 replicate Northern blots. Pixel intensity for each band is normalized to pixel intensity of the 7SL band. Bars represent the average pixel intensity with control set to 1, and dots represent replicate measurements. Error bars represent standard deviation; red asterisk indicates p ≤ 0.05 (n=3-4, ANOVA, Dunnett’s multiple comparisons). B) Heatmap of rRNA processing protein abundance in the control and each of the EXOSC3 variants. Pareto scaling was applied to normalized protein abundance, and clustering uses Euclidean distance and Ward linkage. Colored dots next to protein name indicate proteins that are changing in EXOSC3 variant cells; orange = significantly changing in D132A/D132A cells, purple = G135R/G135R, blue = A139P/A139P (n=4, ANOVA, Tukey HSD).

Several RRPs are increasing in abundance in S1 variant cells, and the colored dots next to each protein indicates the genotypes associated with statistically significant abundance changes (Figure 5B). Increases in proteins associated with rRNA processing suggests the S1 variant cells are likely compensating for decreased RNA exosome abundance and altered rRNA processing through increasing abundance of various rRNA processing proteins. Perhaps these compensation mechanisms are sufficient for rRNA processing rescue in G135R/G135R cells where rRNA processing is relatively unaffected; however, rRNA processing protein upregulation appears insufficient to fully compensate for S1 variant-associated rRNA processing defects since 5.8S rRNA is still significantly decreased by these variants.

### Proteasome inhibition in EXOSC3 S1 variant cells partially rescues RNA exosome abundance

Since we observe most subunit changes in RNA exosome at the protein rather than transcript level, protein complex changes associated with EXOSC3 S1 variants are likely occurring post-translationally. Therefore, we wanted to determine if protein abundance of RNA exosome subunits can be rescued in our S1 variant models by proteasome inhibition. Initial optimization analysis included 10 µM or 25 µM MG132 or DMSO treatment of control cells, and we determined 10 µM MG132 was sufficient to inhibit proteasome activity based on increased abundance of the rapidly turned over proteasome maturation protein (POMP) (Figure S8A).

EXOSC3 control, D132A/D132A and G135R/G135R cell lines were treated with DMSO or 10 µM MG132 for 24 hours, and global protein abundance was analyzed by mass spectrometry (n=3, Figure 6A). Consistent with our western blot analysis, POMP levels significantly increase in 10 µM MG132 treated cells compared to DMSO treated cells indicating successful inhibition of the proteasome in each cell lines (Figure 6B). Protein abundance for each of the RNA exosome subunits was then assessed for a selection of subunits for genotype-treatment interaction analysis using 2-way ANOVA (Figure S8B, Figure 6C-F). EXOSC1 protein levels significantly increased in both EXOSC3 variant cell lines when treated with MG132 compared to DMSO, although overall levels of EXOSC1 remained significantly lower than control cell lines suggesting that MG132 treatment was sufficient to partially rescue EXOSC1 levels (Figure 6C). EXOSC2 protein increased with MG132 treatment in control and EXOSC3 variant cells compared to control, but this was only statistically significant in EXOSC3 G135R/G135R cells (Figure 6C). Quantitative proteomics analysis revealed that EXOSC3 protein abundance was not responsive to 10 µM MG132 for 24 hours in the EXOSC3 variant cells (Figure 6D), indicating that proteasome inhibition was insufficient to rescue EXOSC3 protein levels in the S1 variant cells. Interestingly, we found that EXOSC10 protein was responsive to 10 µM MG132 treatment in both control and variant cell lines compared to DMSO (Figure 6E). The lack of statistical significance between EXOSC10 levels in DMSO treated control cells and MG132 treated variant cells indicates that MG132 treatment is sufficient to recover native EXOSC10 protein abundance levels. Overall, genotype-treatment interaction analysis by 2-way ANOVA revealed no significant interaction for EXOSC1, EXOSC2, EXOSC3, or EXOSC10 (Tables S17-20). No other RNA exosome subunits were significantly rescued by 10 µM MG132 treatment (Figure S8B-C). These data show that MG132 influences all cell lines in a similar manner, with EXOSC3 S1 variant levels being minimally affected by 10 µM MG132 for 24 hours.

**Figure 6:**
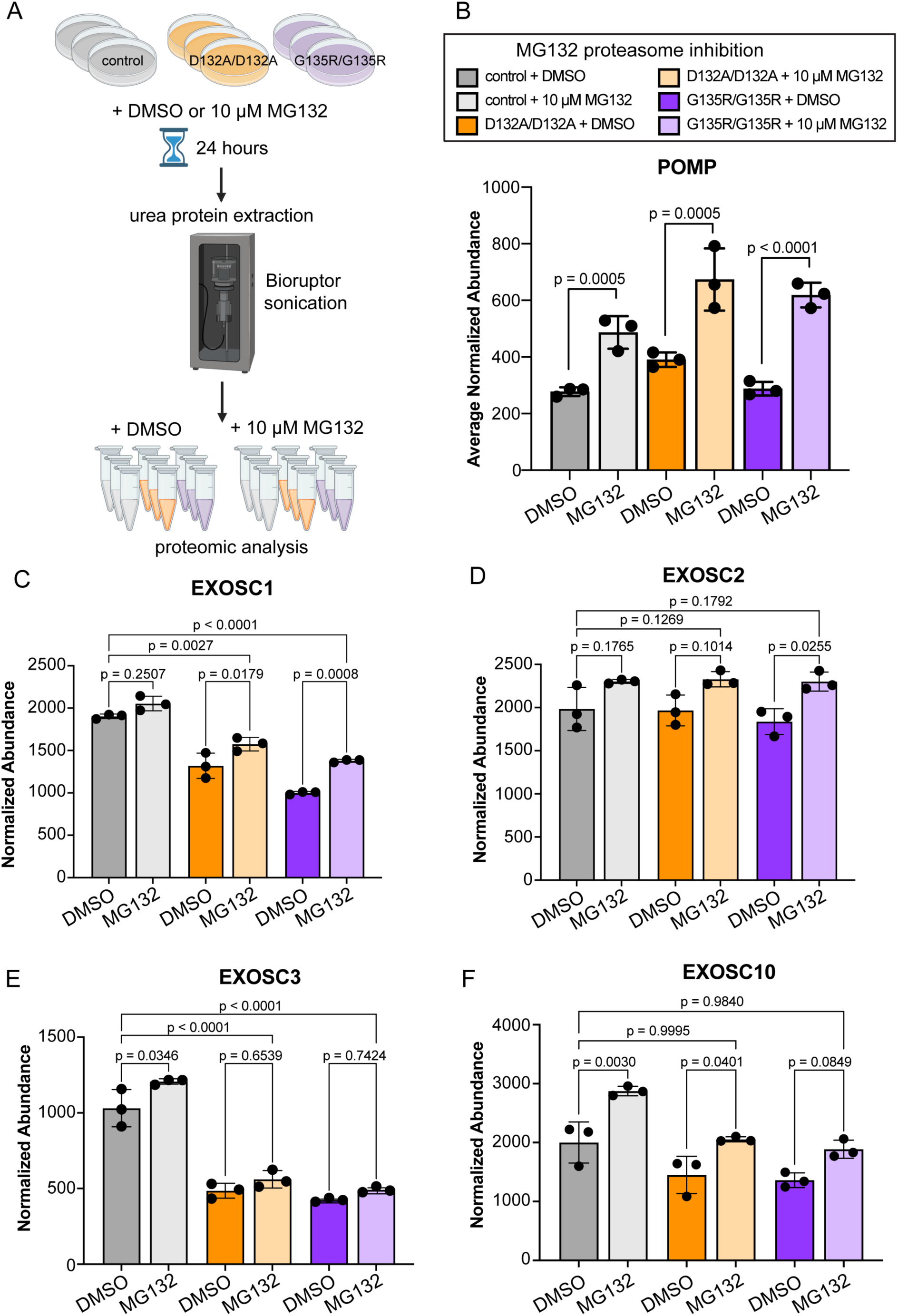
Proteasome inhibition partially rescues RNA exosome subunit abundance in EXOSC3 variants. A) In biological replicates of 3, EXOSC3 control, D132A/D132A and G135R/G135R cells were treated with 10 µM MG132 or DMSO for 24 hours. Proteins were extracted using 8M urea and Bioruptor sonication prior to proteomic analysis. B) Normalized protein abundance of POMP in control and EXOSC3 variant cells treated with DMSO or 10 µM MG132. (ANOVA, Tukey HSD). C-F) Normalized protein abundance of EXOSC1 (C), EXOSC2 (D), EXOSC3 (E), AND EXOSC10 (F) in control and EXOSC3 variant cells treated with DMSO or 10 µM MG132. (2-way ANOVA, Tukey HSD, adjusted p-values shown).

### EXOSC3 overexpression in S1 variant cells rescues RNA exosome subunit abundance

Since we observed multiple changes across the proteome in S1 variant cells, we aimed to determine if overexpression of EXOSC3 leads to system re-equilibration. EXOSC3 D132A/D132A, EXOSC3 G135R/G135R, and control cells were transfected with wild type (WT) EXOSC3 or variant EXOSC3 plasmids (as indicated). Mock transfections were also performed where no DNA was added as negative controls (n=3). After 72 hours, cells were treated with TMR, a fluorescent HaloTag ligand, and sorted based on TMR fluorescence using FACS. TMR positive cells were lysed to extract the whole proteome then analyzed using TMT-based quantitative mass spectrometry to measure protein abundance (Figure 7A). Addition of WT and variant EXOSC3 plasmids to D132A/D132A or G135R/G135R cells significantly increases EXOSC3 protein abundance compared to mock transfected cells (Figure 7B).

**Figure 7:**
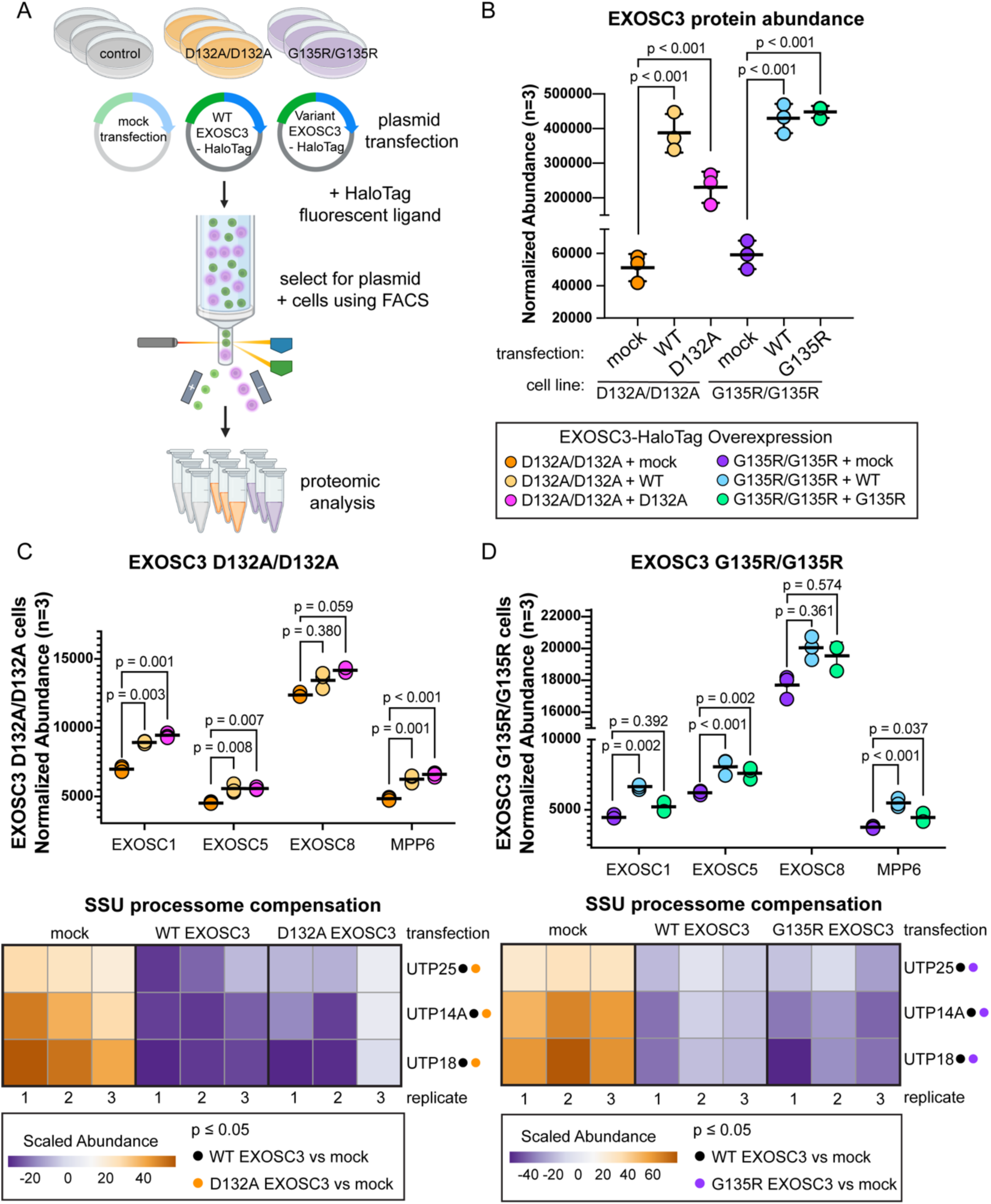
EXOSC3-HaloTag overexpression in EXOSC3 variant cells rescues RNA exosome subunit abundance. A) In biological replicates of 3, control, EXOSC3 D132A/D132A, and EXOSC3 G135R/G135R cells were transfected with a plasmid containing either wild type (WT) or variant EXOSC3-HaloTag cDNA sequence. After 48 hours, cells were treated with a fluorescent HaloTag ligand (TMR), and TMR positive cells were sorted using FACS. Plasmid containing cells were used for proteomic analysis. B) Normalized abundance of EXOSC3 in EXOSC3 D132A and G135R homozygous cells with no DNA, WT EXOSC3-HaloTag, or variant EXOSC3-HaloTag. C, D) Normalized protein abundance of select RNA exosome and SSU processome proteins in D132A/D132A (C) and G135R/G135R (D) cells with no DNA, WT EXOSC3-HaloTag, or variant EXOSC3-HaloTag. Dot plots show protein abundance for EXOSC1, EXOSC5, EXOSC8, and MPP6. P-values were calculated using ANOVA and Tukey HSD post-hoc tests. Heatmaps represent scaled abundance of ribosomal small subunit processome (SSU processome) proteins UTP14A, UTP18, and UTP25. Pareto scaling was applied to normalized protein abundance

Since we observed large changes in protein abundance of subunits interacting in one side of the RNA exosome (Figure 2A – EXOSC1, EXOSC5, EXOSC8, and MPP6), we examined their abundance following EXOSC3 overexpression to determine if increased EXOSC3 could rescue steady-state abundance of other RNA exosome subunits. In D132A/D132A cells, abundance of EXOSC1, EXOSC5, and MPP6 is significantly increased with either WT or D132A EXOSC3-HaloTag overexpression compared to mock transfected cells (Figure 7C). EXOSC8 also increases with both plasmids compared to mock transfection, but did not reach statistical significance (Figure 7C). Similarly, in G135R/G135R cells, WT EXOSC3 overexpression leads to significantly increased EXOSC1, EXOSC5, and MPP6 protein (Figure 7D). Additionally, G135R EXOSC3 overexpression in G135R/G135R cells significantly increased EXOSC5 and MPP6 protein abundance (Figure 7D). These data suggest that WT and variant EXOSC3 protein overexpression is able to interact with other subunits of the RNA exosome resulting in rescue of protein abundance, likely preventing orphan protein decay (58). For G135R/G135R cells in particular, the WT EXOSC3 more robustly increased abundance of the affected subunits, leading us to speculate that the G135R protein less effectively engaged with other RNA exosome subunits relative to WT EXOSC3.

Finally, we wanted to determine if EXOSC3 overexpression resolves other protein abundance changes altered in EXOSC3 S1 variants including factors associated with compensation mechanisms. As in Figure 5, several rRNA processing proteins were significantly increased in abundance in each of the EXOSC3 variant cells, likely to compensate for RNA exosome functional defects seen with reduced efficiency of rRNA processing including production of mature 5.8S rRNA in D132A/D132A cells. We found that overexpression of either WT or D132A EXOSC3 in D132A/D132A cells suppresses abundance of protein components of the ribosomal small subunit processome (SSU processome) including UTP14A, UTP18, and UTP25 (Figure 7C), which were shown to increase in EXOSC3 variant cells relative to control cells (Figure 5). Suppression of SSU processome proteins in G135R/G135R cells overexpressing WT or G135R EXOSC3 also occurred compared to mock controls (Figure 7D). These data show that increased production of EXOSC3 in S1 variant cell lines rescues RNA exosome protein abundance and leads to re-equilibration of RNA processing enzyme abundance outside of the RNA exosome complex.

## DISCUSSION

Our multiomic analyses reveal system-wide responses to EXOSC3 S1 variants including cellular compensation-based changes to respond to changes in abundance and thermal stability of the RNA exosome complex. Proteome changes in the EXOSC3 S1 variant cells support high similarity in impacted cellular processes between both the previously reported pathogenic EXOSC3 variants (D132A and A139P) and the VUS G135R studied here. Comparing to our RNA-seq results, there are more protein abundance changes in EXOSC3 D132A/D132A and EXOSC3 A139P/A139P cells compared to EXOSC3 G135R/G135R cells, however; the overlap of protein level changes between EXOSC3 alleles is much higher than at the RNA level (Figure 1). These findings suggest that each variant has a similar impact on protein complex abundance and stability but downstream functional consequences on RNA related processes are divergent. Considering that our study is the first to directly compare specific PCH1b-associated alleles in human cells, these data are highly significant towards our understanding of the molecular mechanisms that underlie this neurodegenerative disease. As discussed, PCH1b patients experience a range of disease severity in an allele specific manner. Within the RNA exosome, we see that EXOSC3 S1 variants have similar effects on each of the subunits at both the global abundance and stability levels, with DIS3 significantly increasing in each of the variants likely as a compensation mechanism for decreased EXOSC10 stability (Figure 2). The consistent and reproducible increase in DIS3 protein levels could contribute to the greater number of transcripts decreasing in abundance in the EXOSC3 variants. We speculate that in the absence of the fully assembled RNA exosome complex, DIS3 increases could lead to spurious degradation of some RNAs since the targeting and selection mechanisms of the core exosome subunits and associated cofactors for proper RNA targeting is impaired in these S1 variants. We note that protein levels of EXOSC1, EXOSC3, EXOSC5, EXOSC8, and MPP6 (that directly interacts with EXOSC3) are significantly decreased in abundance in all three S1 variants. These subunits are positioned on one face of the RNA exosome complex indicating EXOSC3 D132A, G135R, and A139P variants disrupt steady-state turnover of these subunits. This argument is strengthened by the fact that EXOSC1 levels could be significantly increased by 24-hour treatment with MG132 which inhibits the proteasome.

Although EXOSC10 protein abundance remains relatively unchanged in EXOSC3 S1 variants, it was shown to be significantly destabilized in all three alleles that we tested. The change in EXOSC10 stability is likely associated with the impaired interaction of the exonuclease with the rest of the RNA exosome complex as protein-protein interactions are typically stabilizing (59,60). EXOSC10 directly recognizes and interacts with the SSU processome which is a key step in normal ribosome biogenesis (61,62). Along with EXOSC10, RNA exosome cofactors MPP6 and C1D are directly involved in rRNA maturation (20,21,63,64). Our data highlights the impact of EXOSC3 D132A and A139P variants on rRNA processing, where we see decreased mature 5.8S rRNA (Figure 5A). In all three of the EXOSC3 S1 variants, we note some reduced abundance of MPP6 and C1D with a higher magnitude of thermal destabilization for MPP6, C1D, and EXOSC10 (Figure 2). Thermal destabilization suggests loss of molecular associations between protein complex subunits, and we have previously shown that such changes can be associated with reduced protein complex functional efficiency for the 26S proteasome (59). The RNA data together with the protein data for the EXOSC3 S1 variants also suggests reduced protein complex functional efficiency, with reduced levels of mature rRNAs. We observe an increase in several rRNA processing proteins potentially to compensate for reduced RNA exosome function related to reduced abundance and/or stability of MPP6, C1D, and EXOSC10 (Figure 5B). Interestingly, we do not detect differences in 5.8S rRNA maturation in EXOSC3 G135R/G135R, which could mean the compensation is sufficient to overcome functional changes associated with G135R. Alternatively, the extent of disruption of EXOSC10 interactions with the rest of the complex may be dependent on the precise location of the D132A, G135R, and A139P variants within the RNA exosome complex, allowing for proper EXOSC10 cooperation more of the time. Since these studies were performed in HEK293T cells, it is of note that these RNA processing pathway compensation mechanisms may be less effective in neuronal cells, perhaps specifically Purkinje neurons, leading to cell death as seen in PCH1b.

Overall transcript abundance changes suggest that RNA exosome function is altered to a greater degree in EXOSC3 D132A/D132A cells compared to the other EXOSC3 variants studied (Figures 3A). It is possible that the presence of D132A may specifically disrupt a functional process in the RNA exosome in addition to the protein complex changes that we observed in all S1 variants tested. When analyzing the overlap of transcripts changing in the EXOSC3 S1 variants, we found that EXOSC3 D132A/D132A and EXOSC3 A139P/A139P have the most transcripts in common suggesting these variants lead to more similar changes in RNA exosome functional response (Figure 3D). We also found that rRNA processing is only altered in the pathogenic EXOSC3 variants (Figure 5 and Figure S7) further supporting the stronger impact of EXOSC3 D132A and A139P variants on cellular health compared to the VUS G135R on RNA exosome function. However, these data alone are insufficient to rule out that EXOSC3 G135R variants could cause PCH1b, considering the widespread changes this variant has on RNA exosome protein complex steady-state abundance and stability. Phenotypic characterizations of PCH1b patients have shown that different EXOSC3 missense variants result in varying degrees of disease phenotypes and lifespan expectations (7,8,65–67), suggesting a strong allele-specific effect on functional changes within the RNA exosome. Our data clearly shows that different EXOSC3 variants, even variants that are in the same functional S1 domain and within the same structural loop of the folded protein, have differing impacts on a wide array of downstream cellular events. Careful analyses of EXOSC3 allele specific variant impact on disease model systems is needed to fully understand molecular changes associated with pathology and to predict disease severity. To classify variant pathogenicity, strong evidence is needed linking a variant to disease. The American College of Medical Genetics and Genomics and the Association for Molecular Pathology provide guidelines based on population data, computational predictions, functional studies, and other evidence (4). These studies suggest that wide-spread molecular changes occur in all three EXOSC3 S1 variants, however; it is not yet known what precise threshold of RNA exosome complex dysfunction is sufficient to escape vs. cause PCH1b.

## METHODS

### HEK293T EXOSC3 variant cells

Human embryonic kidney 293 cells with the SV40 T-antigen (HEK293T) were obtained from the American Tissue Culture Collection (ATCC). We chose to use HEK2293T cells for several reasons, one being that the genetic variants of interest are present in all cells within a patient and biophysical changes in proteins will be the same in any cell type. The phenotype of EXOSC3 variants in neuronal cell types typically leads to cell death in patients, and HEK293T cells are more likely to survive despite these variants. HEK293T cells were modified using CRISPR Cas9 technology to introduce missense mutations in the EXOSC3 gene. Guide selection and engineering process were based on previous publications (68,69). Edited cell populations were subjected to single-cell isolation using FACS and then genomic DNA for EXOSC3 was PCR-amplified and screened for introduction of synonymous sequences changes that introduced restriction enzyme sites (BamHI for D132A, BgIII and AatII for G135R, and HindIII for A139P). Positive homozygous and heterozygous clones from this initial screen were further confirmed by EXOSC3 PCR followed by Sanger sequencing.

EXOSC3 variant cell lines used for these experiments include: homozygous EXOSC3 D132A/D132A, EXOSC3 G135R/G135R, EXOSC3 A139P/A139P and heterozygous EXOSC3 +/D132A. A cell line that underwent the same screening procedures as variant cells lines but did not incorporate any of the variants was used as the control cell line. Cells were cultured under 5% CO2-95% air atmosphere at 37°C, in Dulbecco’s modified eagle medium (DMEM) containing 4.5 g/L D-glucose, 10% heat-inactivated fetal bovine serum (FBS), 1X glutaMAX™, and penicillin-streptomycin. All cells were tested with the ATCC mycoplasma test kit and determined mycoplasma-free. All biological replicates are separate growths unless otherwise stated.

### Proteasome inhibition

In biological replicates of 3, control, D132A/D132A, and G135R/G135R cells were plated at 1 million cells per well in a 6-well dish. Cells were treated with 10 µM MG132 or dimethyl sulfoxide (DMSO) vehicle prepared in complete DMEM media. Cells were collected after 24 hours, and pellets were lysed in 8 M urea in 100 mM Tris pH 8.0 and sonicated using a Bioruptor (Diagenode) for 30 cycles (30 s on, 30 s off) on high power.

### Site-Directed Mutagenesis for EXOSC3-HaloTag Overexpression Vector

Site-directed mutagenesis (SDM) was performed using the Agilent QuikChange XL Site-Directed Mutagenesis Kit following the manufacturer’s protocol. Mutagenic primers were designed using the Agilent QuikChange Primer Design tool and synthesized. The mutagenesis reaction was carried out using a plasmid with an N-terminal HaloTag of EXOSC3 on a pFN21A backbone (Promega, FHC10556) as the template. The PCR reaction was prepared with 50 ng template DNA, 125 ng forward primer, 125 ng reverse primer and other reagent amounts according to manufacturer’s protocol. PCR amplification was performed under the following cycling conditions: Initial denaturation: 95°C for 1 min 30 cycles of amplification with Denaturation: 95°C for 50 s, Annealing: 55°C for 1 min, Extension: 68°C for 6 min, Final extension: 68°C for 7 min. Following PCR, 1 µL of DpnI (10 U/µL) was added to the reaction and incubated at 37°C for 1 hour to digest the parental (non-mutated) plasmid. The digested product was then transformed into XL10-Gold ultracompetent cells (Agilent) by heat shock at 42°C for 45 s, followed by incubation on ice for 2 min. Cells were recovered in 500 µL SOC medium at 37°C for 1 hour with shaking at 225 rpm, then plated on LB-agar plates containing ampicillin (100 µg/mL). After overnight incubation at 37°C, individual colonies were selected for mini-prep plasmid isolation. Mutations were confirmed by Sanger sequencing (Genewiz) using appropriate sequencing primers.

### EXOSC3-HaloTag overexpression

Control, D132A/D132A, and G135R/G135R cell lines were plated at a density of 3 million cells per flask. The following day, cells were transfected with 4 µg of plasmid DNA containing wild type or variant EXOSC3-HaloTag plasmid or a no DNA control using polyethylenimine (PEI) transfection reagent. After 72 hours, cells were treated with TMR HaloTag fluorescent ligand, and sorted by flow cytometry. TMR positive cells were retained and pelleted. Cell pellets were lysed in 8 M urea in 100 mM Tris-HCl pH 8.0 and were sonicated using the Bioruptor (Diagenode, Denville, NJ) for 30 cycles (30 s on, 30 s off) on high power.

### Fluorescence-Activated Cell Sorting (FACS) for EXOSC3-HaloTag Analysis

For HaloTag overexpression experiments, cells were incubated with 5 μM tetramethylrhodamine (TMR) HaloTag ligand (MedChemExpress, HY-D2270) in growth media at 37°C for 15 minutes in the dark. Following incubation, cells were washed twice with DPBS and then incubated in fresh growth media for at least 30 minutes before sorting.

For HaloTag overexpression experiments (post-TMR ligand labeling), cells were harvested using TrypLE (Thermo Fisher), washed with DPBS (Dulbecco’s phosphate-buffered saline without magnesium or calcium ions), and resuspended in FACS buffer (DPBS supplemented with 0.5% FBS, 1 mM EDTA, and 1 mM HEPES). The cell concentration was adjusted to 7–8 × 10⁶ cells/mL, and samples were passed through a 40 μm cell strainer to remove aggregates.

FACS was performed using a Sony MA900 cell sorter equipped with 488 nm and 561 nm laser lines and corresponding emission filters (FL2 585/30 for TMR). Data acquisition and sorting were conducted using the Sony MA900 Cell Analyzer software (version 3.1.1). Sorting gates were defined based on forward scatter (FSC), back scatter (BSC), and marker expression (TMR intensity for HaloTag overexpression). Cells were selected based on high FSC-A and BSC-A, with doublets excluded using FSC-H vs. FSC-A gating. After gating, cells with high marker intensities were collected. Target populations were sorted into DPBS using the “Normal” sorting setting. Approximately, 700,000 to 1,200,000 marker positive cells were collected.

Sorted cells were centrifuged at 1000 × g for 5 minutes at 4°C and resuspended in 8 M urea in 100 mM Tris-HCl (pH 8.5). Samples were either stored at -80°C for future processing or sonicated using the Bioruptor (Diagenode, Denville, NJ) for 30 cycles (30 s on, 30 s off) on high power. Samples were normalized to 50 μg of protein in 100 μL in 8 M Urea in 100 mM Tris-HCl buffer and processed for mass-spectrometry-based proteomics analysis.

### Sample collection for global proteomics

In replicates of 4, approximately 10 million cells were collected per genotype. Proteins were extracted by denaturing with 8M urea in 100 mM Tris (pH 8.5). Samples underwent sonication using a Bioruptor (Diagenode, Denville, NJ) for 30 cycles with 30 seconds on, 30 seconds off at high power. Samples were centrifuged at 12,000 rcf for 20 minutes at 4°C. 1 µL of benzonase (0.1 unit/µL) was added to each sample and incubated for 30 minutes at 37°C with vortexing throughout. Samples were centrifuged at 12,000 rcf for 20 minutes at 4°C, and supernatant was reserved.

### Sample collection and heat treatment for PISA and TPP

In replicates of 4, approximately 40 million cells were collected per genotype (control, D132A/D132A, G135R/G135R, A139P/A139P, P/D132A, P/G135R). Cells were resuspended in NP-40 lysis buffer and lysed by 3 freeze thaw cycles in liquid nitrogen (30 s in liquid nitrogen, 10 minutes at 4°C). Following centrifugation at 20,000 rcf for 30 min, protein concentrations were determined using a detergent compatible protein assay kit. Samples were adjusted to the same concentration and divided equally into 12 tubes for heat treatment. Samples underwent heat treatment in the thermocycler (25°C for 3 minutes, 3 minutes at temperature gradient, 25°C for 3 minutes) (Figure 4). Temperature gradient is as follows: 25°C, 35°C, 39.3°C, 43.3°C, 48.6°C, 50.1°C, 51.9°C, 54.5°C, 58.7°C, 60.6°C, 74.9°C, 90°C. Each sample was centrifuged to pellet aggregated protein, and supernatant was reserved. For PISA, 10 uL of the following temperature treated samples were combined: 39.3°C, 43.3°C, 48.6°C, 50.1°C, 51.9°C, 54.5°C, 58.7°C, 60.6°C (Figure 5). Proteins were precipitated in 20% trichloroacetic acid (TCA) overnight and washed with acetone. Precipitated proteins were resuspended in 8M Urea in 100 mM Tris-HCl (pH 8.0).

### Reduction, alkylation, and digestion

Proteins were reduced with 5 mM tris(2-carboxyethyl)phosphine (TCEP) and alkylated with 10 mM chloroacetaminde (CAM). Samples incubated with trypsin/Lys-C mix (1:70 protease:substrate ratio) for 4 hours, then were diluted with 100 mM Tris-HCl to reduce urea concentration to less than 1M and digested overnight. Digestion was quenched with 0.4% trifluoroacetic acid (TFA). Peptides were desalted on a 50 mg Sep-Pak C18 Vac (Waters Corporation) using a vacuum manifold. Peptides were eluted with 70% acetonitrile (ACN) and 0.1% formic acid (FA), dried by speed vacuum, and resuspended in 50 mM triethylammonium bicarbonate (TEAB, pH 8.5).

### TMT labeling

Equal volumes of each sample were labeled with tandem mass tags (TMT) for 2 hours at room temperature. TMTpro 16 or 18plex sets were used for all RNA exosome experiments.

Labeling reactions were quenched with 5% hydroxylamine, and labeled peptides were mixed together. TMT lot numbers and labeling scheme can be provided upon request.

### High pH reversed-phase fractionation

Peptide mixtures were desalted and fractionated using 50 mg Sep-Pak C18 Vac columns (Waters Corporation). Peptides were eluted in 12.5%, 15%, 17.5%, 20%, 22.5%, 25%, 30%, and 70% acetonitrile in 0.1% triethylamine. Fractions were dried in speed vacuum and resuspended in 0.1% formic acid.

### Data acquisition

Nano-LC–MS/MS analyses were performed using an Orbitrap Eclipse mass spectrometer (Thermo Fisher Scientific) coupled to an EASY-nLC high-performance liquid chromatography (HPLC) system (Thermo Fisher Scientific). Each fraction was loaded onto an Aurora Series UHPLC C18 1.7 µm column (IonOpticks, AUR3-25075C18A, 25 cm x 75 µm). HPLC and mass spectrometer settings for each experiment are available upon request. For global proteomics and PISA experiments, two technical replicates, meaning the same samples were run on the mass spectrometer twice, were obtained.

### Protein identification and quantification

Resulting MS^2^ RAW files were analyzed in Proteome Discoverer™ 2.5 (Thermo Fisher Scientific) with FASTA databases including UniProt human sequences plus common contaminants. FASTA database used for RNA exosome experiments also contained variant protein sequences of EXOSC3. Quantification methods used isotopic impurity levels available from Thermo Fisher Scientific. SEQUEST HT searches were conducted with a maximum number of two missed cleavages, a precursor mass tolerance of 10 parts per million, and a fragment mass tolerance of 0.02 Da. Static modifications used for the search were (i) carbamidomethylation on cysteine (C) residues and (ii) TMT or TMTpro labels on lysine (K) residues and the N termini of peptides. Dynamic modifications used for the search were oxidation of methionine, acetylation of N termini, and N-terminal loss of methionine with and without acetylation. Percolator false discovery rate (FDR) was set to a strict setting of 0.01 and a relaxed setting of 0.05. Values from both unique and razor peptides were used for quantification. For global and PISA experiments, data was normalized by total peptide amount without scaling. For the proteasome inhibition experiment, proteins were normalized to 28 of the most abundant proteins that were not impacted by MG132 treatment. Overexpression data was normalized to total peptide amount of all proteins except for EXOSC3.

### RNA extraction

For RNA sequencing, approximately 10 million cells were grown as described above and collected in 1 mL TRIzol™ reagent (Life Technologies, Carlsbad, CA). The Direct-zol RNA mini prep kit (Zymo Research, Orange, CA) was used to extract RNA. 100 µL of collected cells in TRIzol™ reagent was added to 300 µL of TRIzol to dilute samples 1:4. Manufacturer protocol was followed for remaining RNA extraction steps. Isolated RNA concentration was measured using a NanoDrop 2000 Spectrophotometer (Thermo Scientific, Waltham, MA).

For northern blot analysis, 200 µL of collected cells in TRIzol™ reagent were combined with 800 µL TRIzol™ reagent. Samples were pipetted 10 times to ensure no clumps of cells remained. These incubated at room temperature for 5 minutes. 200 µL of chloroform was added and incubated for 3 minutes at room temperature. Following centrifugation at 20,000 rcf for 15 minutes at 4°C, the samples separated into 3 layers. The top layer containing RNA was moved to a new tube. 500 µL isopropanol was added to RNA, mixed by inversion, and incubated at room temperature for 10 minutes. The samples were centrifuged at 20,000 rcf for 10 minutes at 4°C and the supernatant was removed. The pellet was washed with 1mL 80% ethanol and centrifuged at 7,500 rcf for 5 minutes at 4°C. The supernatant was removed, and the pellet air dried for approximately 10 minutes. RNA pellets were resuspended in molecular biology grade water and incubated at 56°C for 10 minutes. RNA quality was checked using the Agilent TapeStation, and a RIN of 8 or higher was required to move to northern blot analysis.

### Library preparation and sequencing

Total RNA samples were first evaluated for the quantity and quality using Agilent TapeStation. All the samples had good quality with RIN (RNA Integrity Number) greater than 8. 100 nanograms of total RNA was used for library preparation with the KAPA total RNA Hyperprep Kit (KK8581) (Roche). ERCC mixture of synthetic RNAs was spiked in every total RNA sample. Each resulting uniquely dual-indexed library was quantified and quality accessed by Qubit and Agilent TapeStation; Multiple libraries were pooled in equal molarity. The pooled libraries were sequenced on an Illumina NovaSeq 6000 sequencer. Around 200 million 100 bp paired-end reads were generated for each sample. RNA sequencing data was uploaded to Gene Expression Omnibus (GEO) Accession XXXX.

### RNA sequencing data processing

The FastQ files from the RNA sequencing were processed based on the following pipeline. First, adapter sequences were trimmed using bbduk tool from BBTools v.38.72 (70)with the following parameters: *k=23 ktrim=r hdist=1 tpe=t tbo=t useshortkmers=t mink=11 qtrim=rl trimq=10 minlength=50*. Reference FASTA file of adapters (*adapter.fa* from the BBTools suite). FastQC v. 0.11.5 was used to determine the quality of trimmed reads (71).

Before indexing, a FASTA reference genome was made by concatenating the Ensembl import GRCh38 version 108 with the ERCC Spike-In sequences. An indexed human reference genome with ERCC Spike-In was made using STAR v.2.7.3a with parameters *-sjdbOverhang 100* (72). The trimmed reads were aligned to the indexed reference genome using STAR v. 2.7.3a using the parameters *--outFilterType BySJout --outFilterMultimapNmax 20 --alignSJoverhangMin 8 -- alignSJDBoverhangMin 1 --outFilterMismatchNmax 999 --outFilterMismatchNoverLmax 0.04 --alignIntronMin 20 --alignIntronMax 1000000 --alignMatesGapMax 1000000 --outSAMattributes NH HI NM MD --outSAMtype BAM SortedByCoordinate* (73). Gene read counts and junction spanning reads were obtained using the featureCounts tool of Subread v. 2.0.3 with the following parameters accounting for the reverse stranded nature of the library generation: -*p -- countReadPairs -s 2* and GTF used in STAR genome indexing previously mentioned (74).

### Differential Gene Expression Analysis

The gene read count files were imported into RStudio running R v.4.2.3. Differential expression analysis was performed using DESeq2 v. 1.38.3 (75). DESeq data set was created using the *DESeqDataSetFromMatrix* function with the design argument *∼ Genotype* (the EXOSC3 genotype) and the reference level to EXOSC3 control genotype. Only genes with total read counts greater than or equal to 5 were included in the estimations of size factors, dispersion, and negative binomial GLM fitting. In the estimation of size factors, a vector of counts aligned to the ERCC Spike-In was used as the *controlGenes* argument in the function *estimateSizeFactors*. A log-fold-change shrink was applied using “*ashr*” (76). Differential expressed genes were considered to be genes with an FDR-adjusted (Benjamini-Hochberg) *p*-value less than or equal to 0.05. Data is provided in Tables S11-13.

### Enrichment analysis of untranslated region (UTR) elements in differentially abundant transcripts

To assess the overrepresentation of known UTR elements in differentially abundant transcripts, we obtained manually annotated regulatory sites for trans-factors and cis-elements in UTRs from the Atlas of UTR Regulatory Activity (AURA) version 2.6, filtering for human-specific annotations (57,56). Overrepresentation analysis for each regulatory element was performed using Fisher’s exact test, implemented in the R package *stats*, with p-values adjusted using the Benjamini-Hochberg correction. The top 20 most statistically significant regulatory elements were visualized using *ggplot2*.

### Northern blot

RNAs were separated using a 1% agarose/formaldehyde gel and passively transferred by capillary overnight to a Hi-bond nylon membrane. Membranes were probed using toligonucleotide probes against their indicated targets. Bands were quantified using Image Lab software (Bio-Rad).

### Western blot

Samples were lysed in RIPA buffer and equal volumes of sample were used then normalized to beta-actin intensity. Proteins were separated using a BioRad Mini-Protean TGX stain free precast gel or Criterion TGX anykD precast gel and transferred to PVDF membrane by electrophoresis. Primary antibodies used include rabbit anti-EXOSC3 (1:5000 dilution; Protein Tech #15062-1-AP), mouse anti-beta actin (1:5000 dilution; Sigma #0000120485), rabbit anti-POMP (1:1000 dilution; Abcam #ab170865). Secondary antibodies used include HRP conjugated Goat anti-rabbit IgG (1:5000 dilution; BioRad #1706515) and HRC conjugated Goat anti-mouse IgG (1:5000; BioRad #1706516). Clarity Western ECL substrate (BioRad) was used to develop the HRP signal, and membranes were imaged using BioRad ChemiDoc Imager.

### Gene ontology

Gene ontology (GO) analysis was performed using ShinyGo version 8.0(77). KEGG and biological process enrichments were used for these experiments. Protein-protein interaction analysis was performed using STRING with high confidence (0.700) (78–82).

### Data visualization

Volcano plots were created in R Studio using ggplot2 (version 3.4.2) (83). UpSet plots were created in R studio using UpSetR (84). Bar plots were created in GraphPad Prism 10.

Venn diagrams were generated using Venny (version 2.1) (85). Heatmaps were generated using ClustVis (86). 2-way ANOVA and Tukey analysis were performed using GraphPad Prism 10 and data are in Tables S17-20.

## Funding

R01NS121550 and P30CA082709 to A.L.M., R35GM146888 to J.Z.V., R35GM138123 to H.G., T32CA2723370 to A.M.R., F30AG079580 and T32GM148382 to H.R.S.W.

## Author’s Contributions

A.M.R. – experimental design, data acquisition, data analysis, writing and revision of manuscript

H.R.S.W. - experimental design, data acquisition, data analysis, writing and revision of manuscript

M.P.B. – data analysis, writing and revision of manuscript

W.R.S.K. – data acquisition

J.D.R. – experimental optimization

S.A.P.J. – writing and revision of manuscript

L.A.C. – data acquisition

A.H. – data acquisition

H.G. – data acquisition

S.P. – cell line generation

E.H.D. – data acquisition

J.Z.V. – funding acquisition, supervision, experimental design, data analysis, revision of manuscript

A.L.M. – funding acquisition, project administration, experimental design, data analysis, writing and revision of manuscript

## Acknowledgements

Cell lines were generated in collaboration with the Indiana University School of Medicine Gene Editing Core. The RNA sequencing work was performed in collaboration with the Indiana University School of Medicine Center for Medical Genetics. We thank Yunlong Liu and Rudong Li for their assistance with data analysis. The mass spectrometry work performed in this work was done by the Indiana University School of Medicine Center for Proteome Analysis.

Acquisition of the IUSM Center for Proteome Analysis’ instrumentation used for this project was provided by the Indiana University Precision Health Initiative. The Center for Proteome Analysis receives support from 3UL1TR002529 and P30CA082709. The authors acknowledge the Indiana University Pervasive Technology Institute for providing supercomputing and storage resources that have contributed to the research results reported within this paper.

---

**Figure S1:**
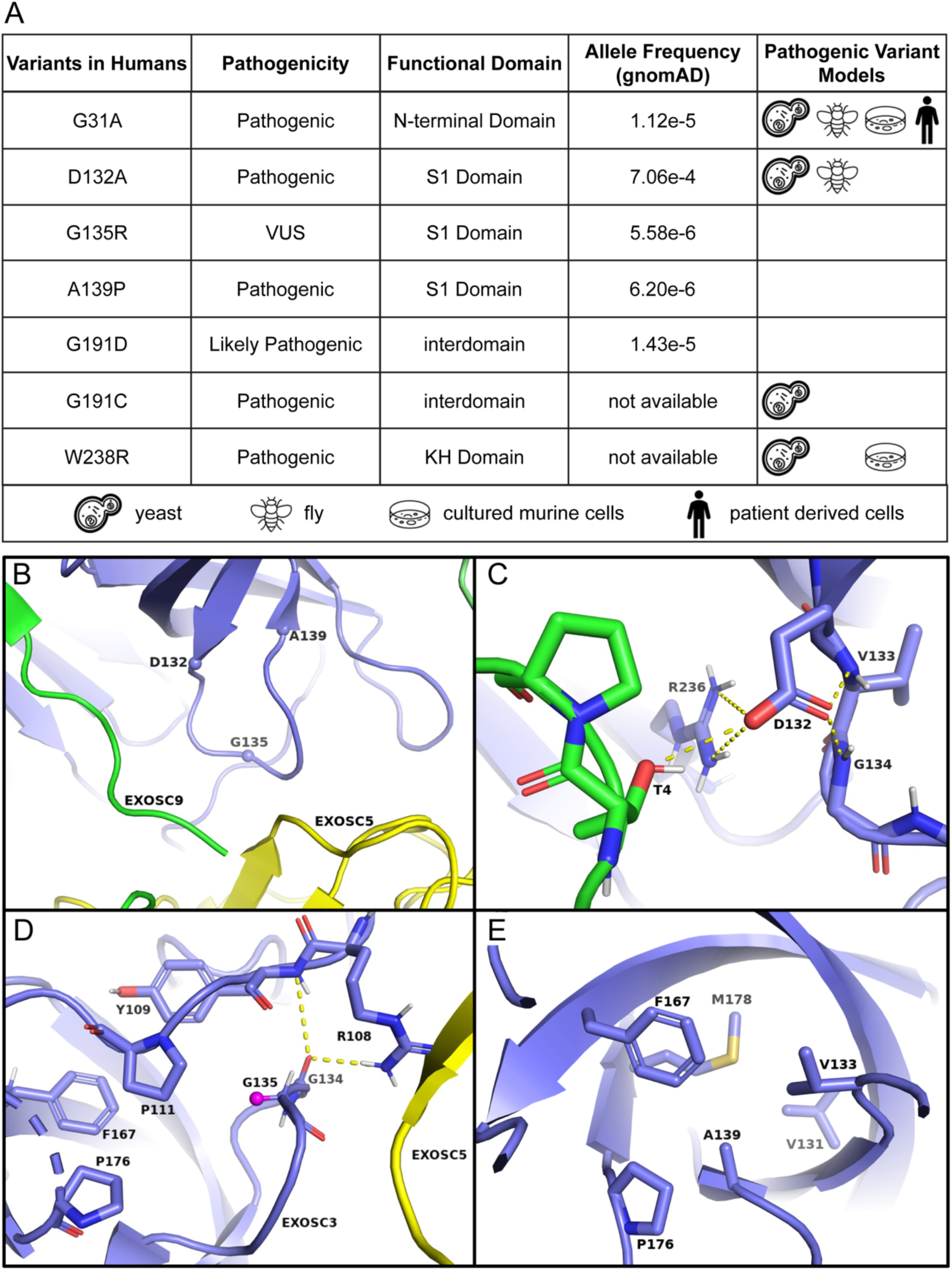
EXOSC3 variants associated with PCH1b. A) Table of EXOSC3 variants and current models used for study. VUS stands for variant of uncertain significance, S1 and KH domains are RNA binding domains, and the N-terminal domain is important for structure and intersubunit interactions. B) A zoomed in EXOSC3 structural image is shown in blue with D132, G135, and A139 residues represented as spheres (PDBID 6D6Q) (43). Surrounding RNA exosome subunits EXOSC9 and EXOSC5 are shown in green and yellow, respectively. C) D132 interacts with residues in EXOSC3 and EXOSC9. D) G135R is in close proximity to residues in EXOSC5 and EXOSC3. Glycine’s second alpha hydrogen that is replaced with a L-amino acid side chain is shown as a pink sphere. E) A139P is in close proximity to other EXOSC3 residues in the S1 beta-barrel structure. Dashed yellow lines represent hydrogen bonding.

**Figure S2:**
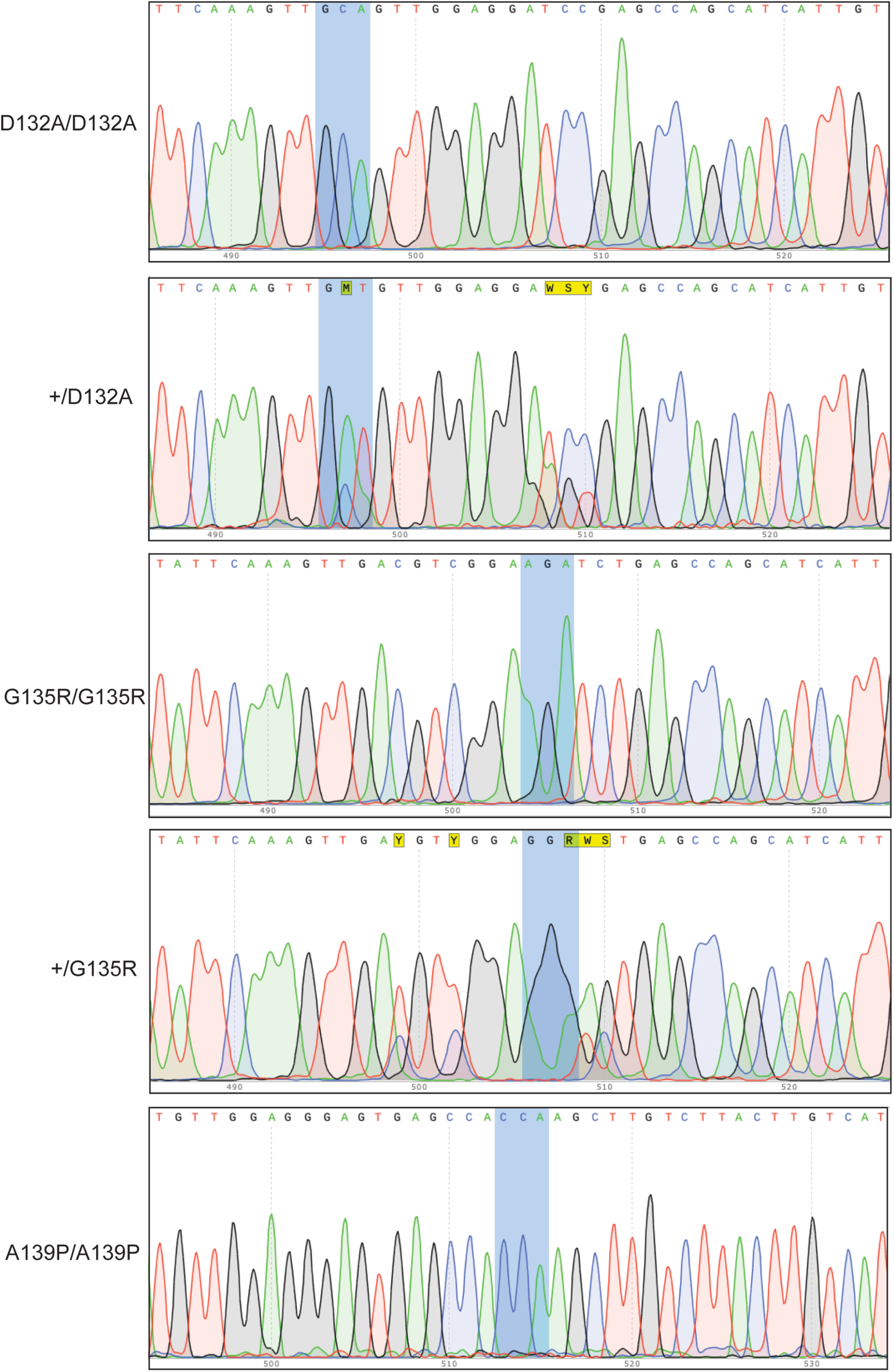
Sanger sequencing confirmation of CRISPR-Cas9 generated EXOSC3 variant cell lines. Blue boxes indicate nucleotide changes.

**Figure S3:**
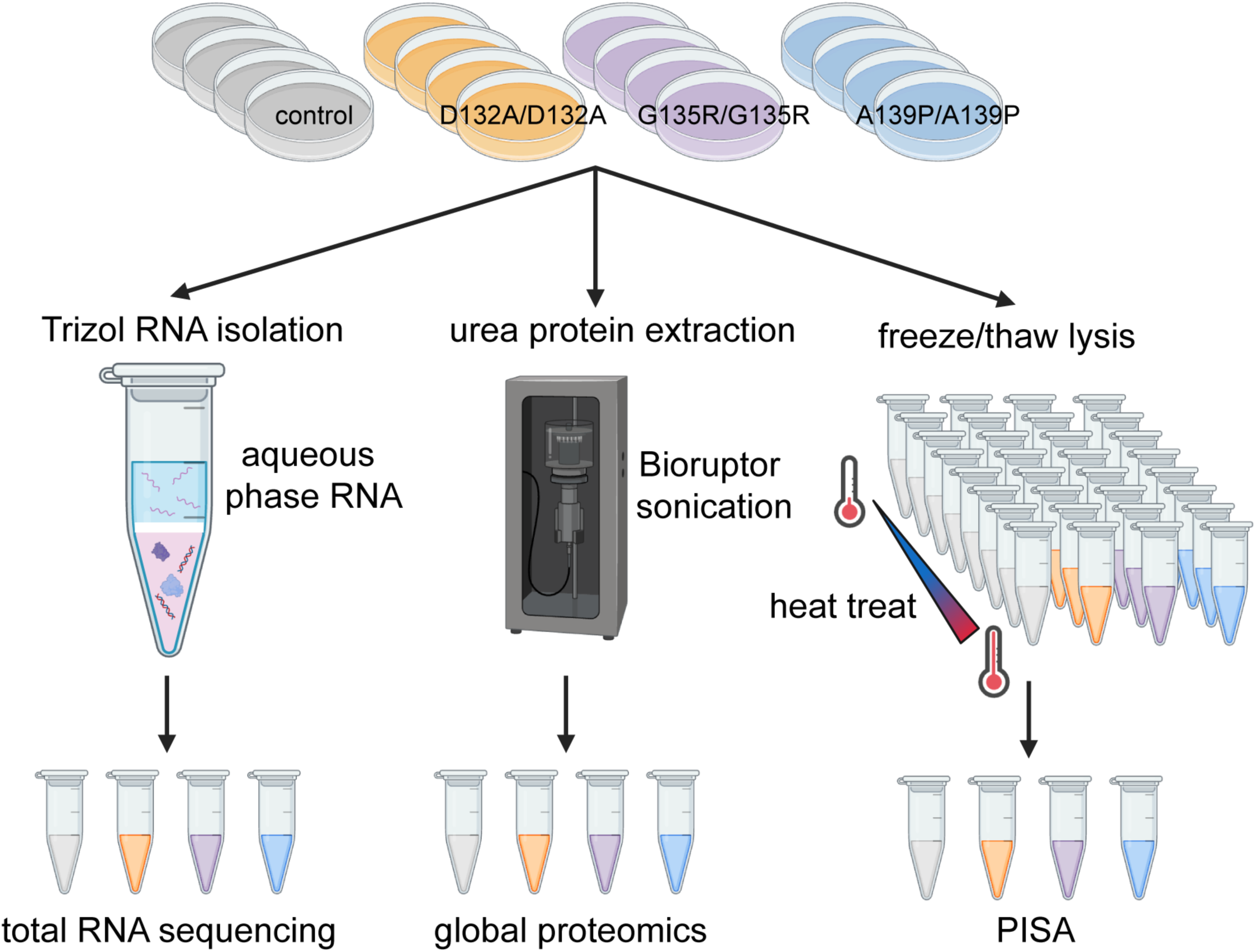
Experimental workflow. Control and EXOSC3 variant cells were grown in biological replicates of 4. Cells were used for RNA isolation, protein extraction, and PISA.

**Figure S4:**
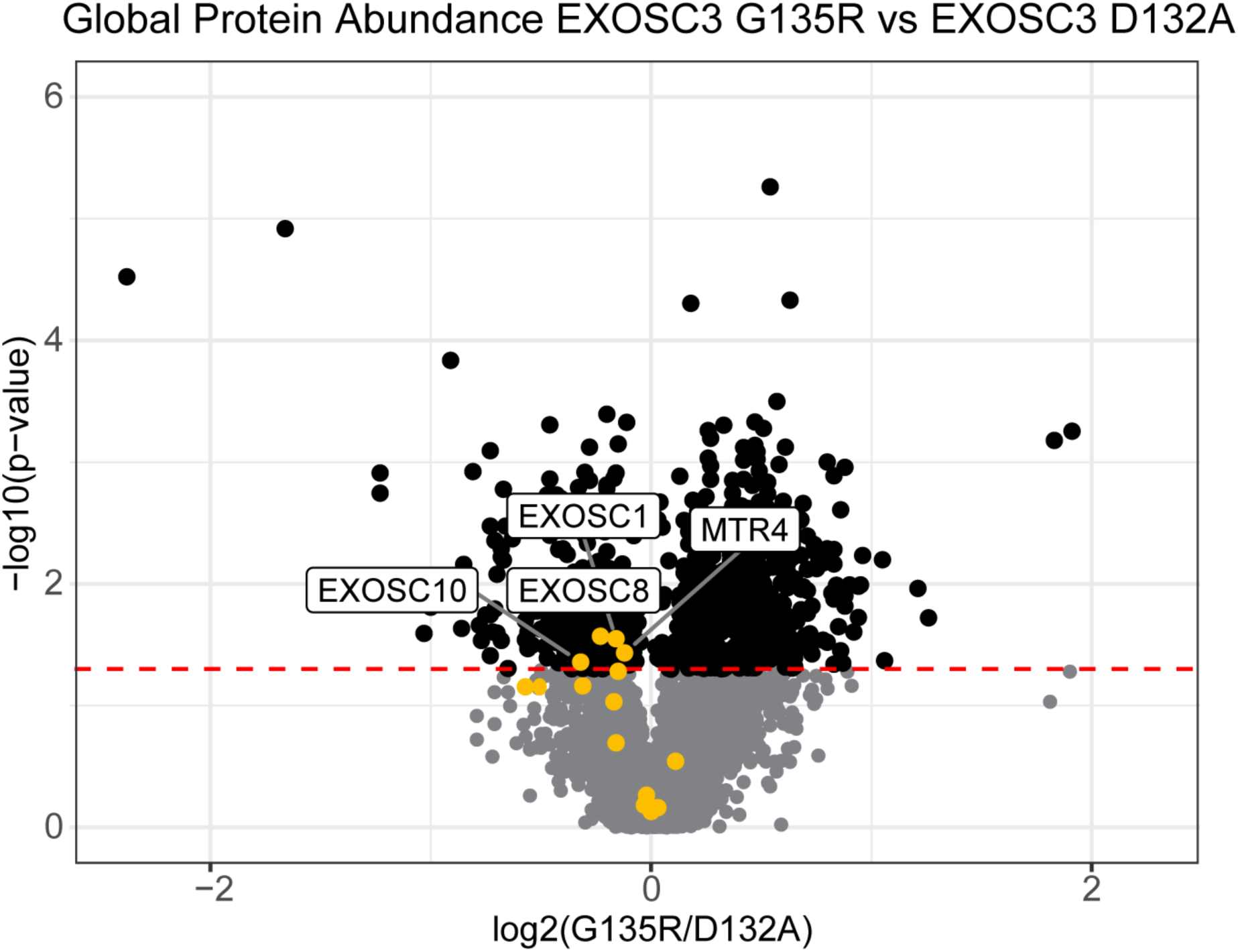
Protein abundance changes in G135R/G135R cells compared to D132A/D132A cells. Points in yellow represent RNA exosome subunits.

**Figure S5:**
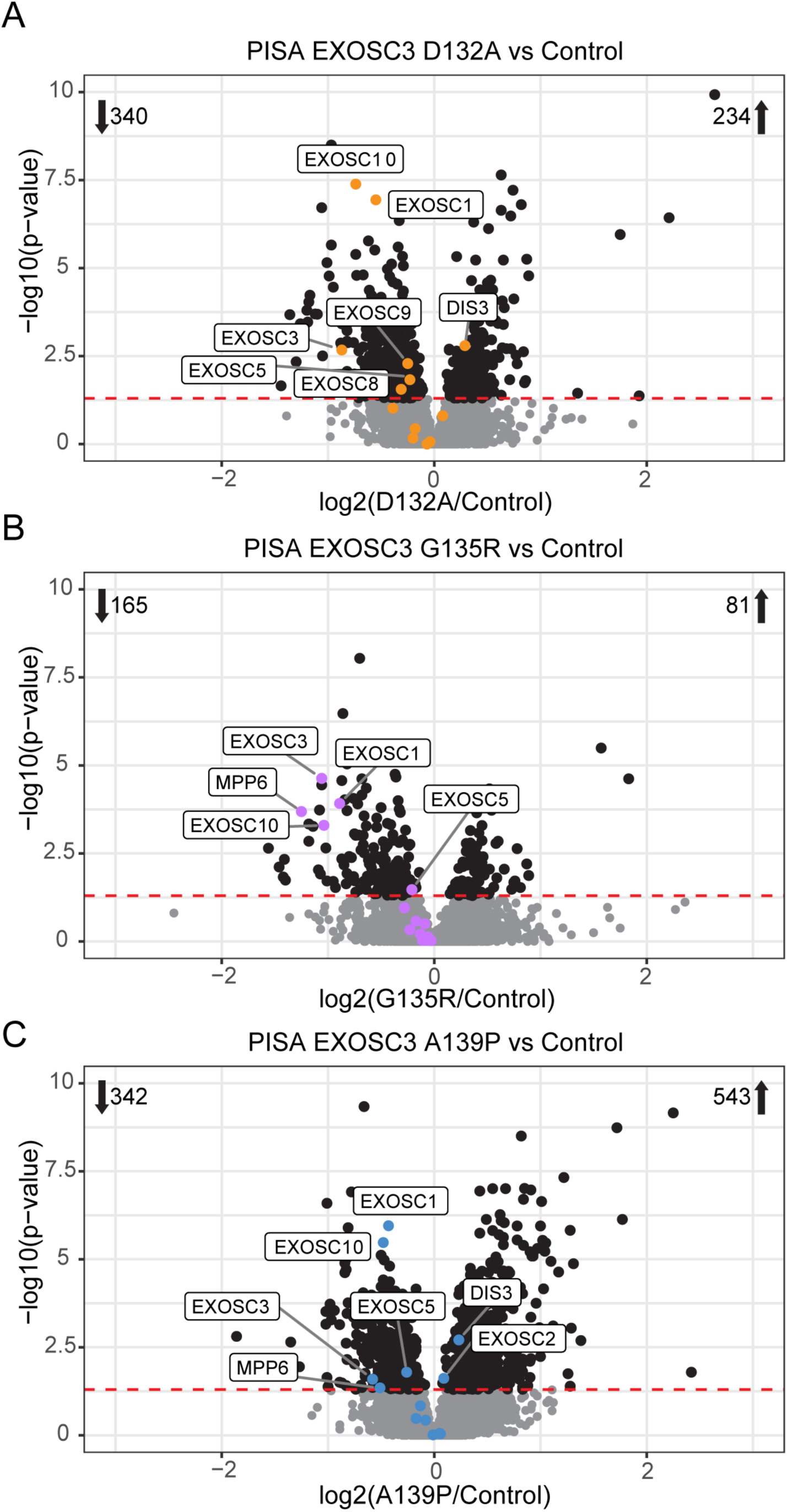
Changes in protein thermal stability identified by PISA. A) EXOSC3 D132A/D132A cells compared to control. B) EXOSC3 G135R/G135R cells compared to control. C) EXOSC3 A139P/A139P cells compared to control. The log2(fold change) for each protein was plotted against the -log10 of the p-value. The red dashed line represents p-value cut off of 0.05. Each point represents one protein, and black points represent significantly changing proteins. Orange, purple, and blue points represent RNA exosome subunits. Number of proteins significantly increasing or decreasing in stability are shown in the corners of each graph.

**Figure S6:**
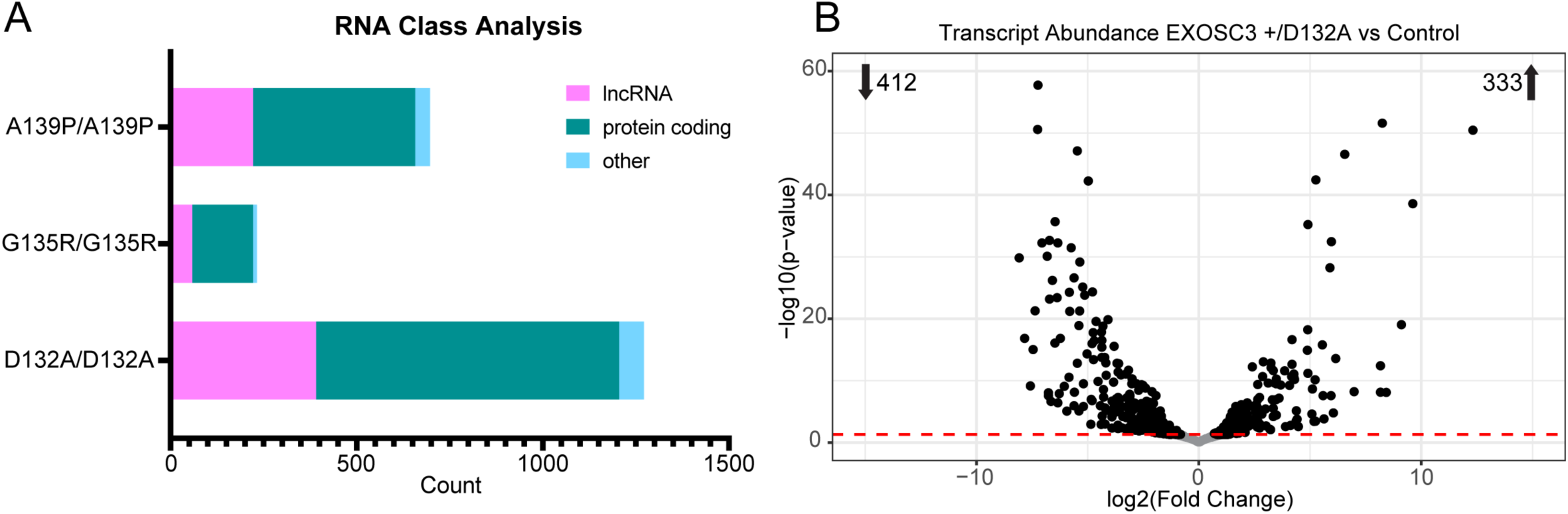
Transcript changes in EXOSC3 variant cells. A) Types of RNAs significantly changing in each of the EXOSC3 variants. Count of differentially abundant transcripts is plotted for each genotype. Different color bars represent different types of RNA. Stacked bars add up to the total number of RNAs significantly changing. B) EXOSC3 +/D132A compared to control. The log2(fold change) for each transcript was plotted against the -log10 of the adjusted p-value. Each point represents one transcript. The red dashed line represents an adjusted p-value cut off value of 0.05. Number of transcripts significantly increasing or decreasing are shown in the corners of each graph.

**Figure S7:**
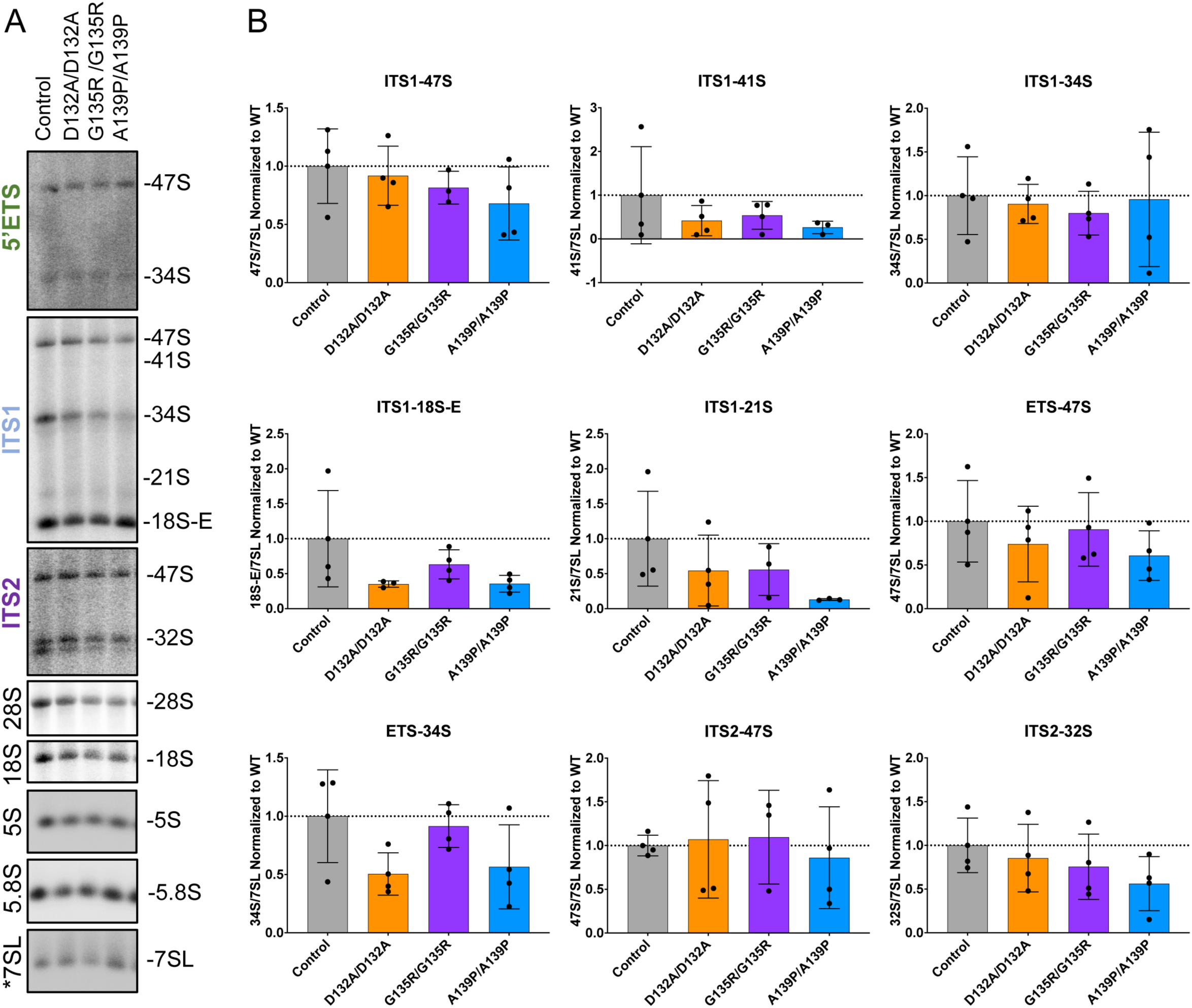
Northern blot analysis of mature and intermediate rRNA abundance levels (A). One representative replicate of EXOSC3 variants and control is shown. Labels on the left indicate which probe was used and identify the precursor or mature rRNA detected. B) Quantification of premature rRNAs from 4 replicates. Pixel intensity for each band is normalized to pixel intensity of the 7SL band. Bars represent the average pixel intensity with control set to 1, and each dot represents the replicates. Error bars represent standard deviation.

**Figure S8:**
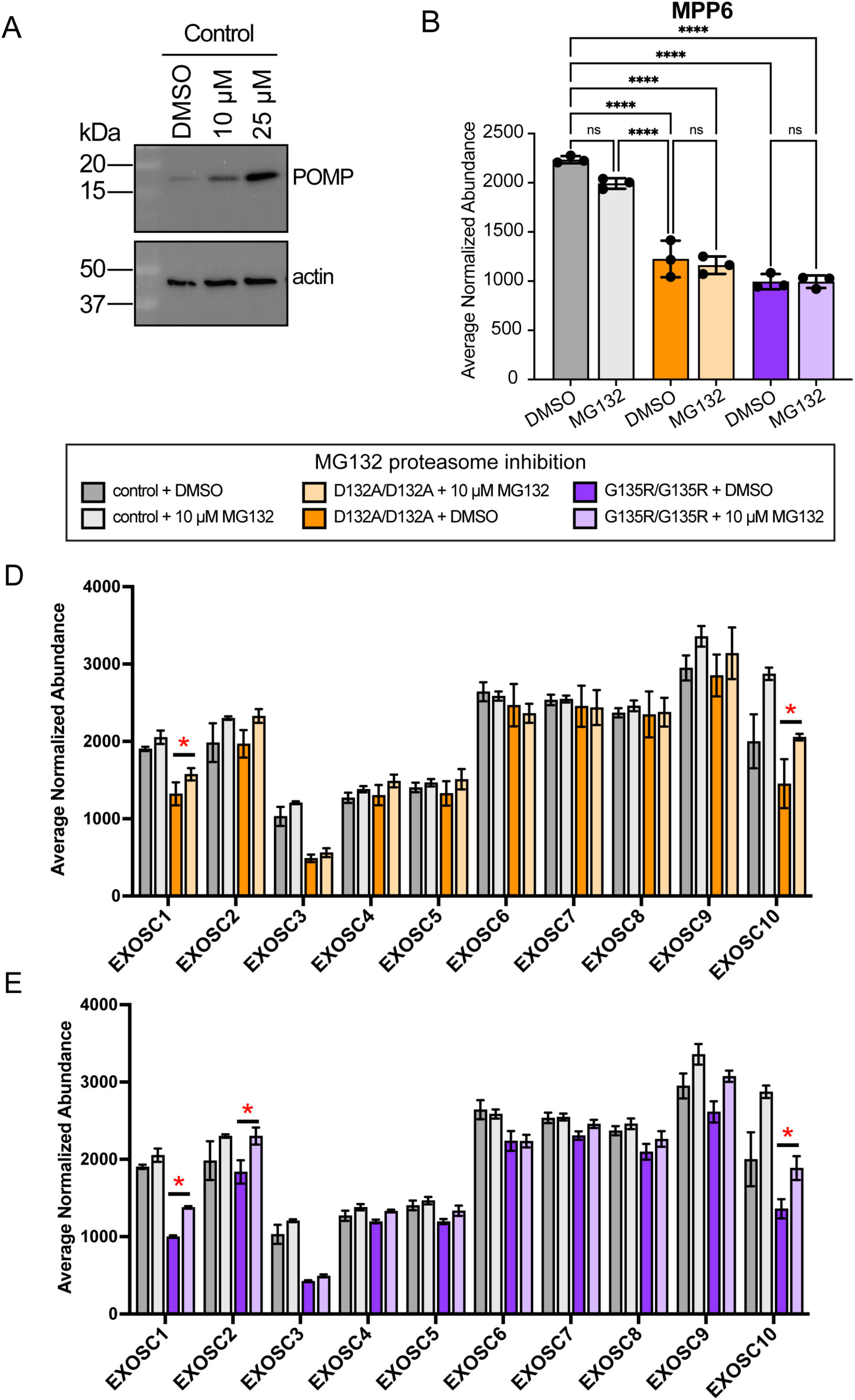
Proteasome inhibition partially rescues RNA exosome subunit abundance in EXOSC3 variants. A) Western blot analysis of POMP, EXOSC3, and actin (loading control) in control and EXOSC3 D132A/D132A cells treated with 10 µM MG132 or DMSO. B) Normalized protein abundance of MPP6, a cofactor of the RNA exosome, is shown for each of the cell lines. Statistical significance is represented with asterisks (2-way ANOVA, Tukey HSD, adjusted p ≤ 0.05). C, D). Normalized protein abundance of RNA exosome subunits in control, EXOSC3 D132A/D132A (C), and G135R/G135R (D) cells treated with DMSO or 10 µM MG132.

## Notes

### Competing Interest Statement

The authors have declared no competing interest.

## References

1. Chin L, Hahn WC, Getz G, Meyerson M. Making sense of cancer genomic data. Genes Dev. 2011 Mar 15;25(6):534–55.

2. Hutter C, Zenklusen JC. The Cancer Genome Atlas: Creating Lasting Value beyond Its Data. Cell. 2018 Apr 5;173(2):283–5.

3. Amberger J, Bocchini C, Hamosh A. A new face and new challenges for Online Mendelian Inheritance in Man (OMIM®). Hum Mutat. 2011 May;32(5):564–7.

4. Richards S, Aziz N, Bale S, Bick D, Das S, Gastier-Foster J, et al. Standards and Guidelines for the Interpretation of Sequence Variants: A Joint Consensus Recommendation of the American College of Medical Genetics and Genomics and the Association for Molecular Pathology. Genet Med Off J Am Coll Med Genet. 2015 May;17(5):405–24.

5. Morton DJ, Kuiper EG, Jones SK, Leung SW, Corbett AH, Fasken MB. The RNA exosome and RNA exosome-linked disease. RNA. 2018 Feb;24(2):127–42.

6. Barth PG. Pontocerebellar hypoplasias. An overview of a group of inherited neurodegenerative disorders with fetal onset. Brain Dev. 1993 Dec;15(6):411–22.

7. Eggens VR, Barth PG, Niermeijer JMF, Berg JN, Darin N, Dixit A, et al. EXOSC3 mutations in pontocerebellar hypoplasia type 1: novel mutations and genotype-phenotype correlations. Orphanet J Rare Dis. 2014 Dec;9(1):1–10.

8. Rudnik-Schöneborn S, Senderek J, Jen JC, Houge G, Seeman P, Puchmajerová A, et al. Pontocerebellar hypoplasia type 1. Neurology. 2013 Jan 29;80(5):438–46.

9. Wan J, Yourshaw M, Mamsa H, Rudnik-Schöneborn S, Menezes MP, Hong JE, et al. Mutations in the RNA exosome component gene EXOSC3 cause pontocerebellar hypoplasia and spinal motor neuron degeneration. Nat Genet. 2012 Jun;44(6):704– 8.

10. Schwabova J, Brozkova DS, Petrak B, Mojzisova M, Pavlickova K, Haberlova J, et al. Homozygous EXOSC3 mutation c.92G→C, p.G31A is a founder mutation causing severe pontocerebellar hypoplasia type 1 among the Czech Roma. J Neurogenet. 2013 Dec;27(4):163–9.

11. Schottmann G, Picker-Minh S, Schwarz JM, Gill E, Rodenburg RJT, Stenzel W, et al. Recessive mutation in EXOSC3 associates with mitochondrial dysfunction and pontocerebellar hypoplasia. Mitochondrion. 2017 Nov;37:46–54.

12. Biancheri R, Cassandrini D, Pinto F, Trovato R, Di Rocco M, Mirabelli-Badenier M, et al. EXOSC3 mutations in isolated cerebellar hypoplasia and spinal anterior horn involvement. J Neurol. 2013 Jul;260(7):1866–70.

13. Szeto CH, Rubin S, Sidlow R. Homozygous EXOSC3 c.395A>C Variants in Pontocerebellar Hypoplasia Type 1B: A Sibling Pair With Childhood Lethal Presentation and Literature Review. Cureus. 2023 May;15(5):e39226.

14. EXOSC3 | gnomAD v4.1.0 | gnomAD [Internet]. [cited 2024 May 29]. Available from: https://gnomad.broadinstitute.org/gene/ENSG00000107371?dataset=gnomad_r4

15. Fox MJ, Mosley AL. Rrp6: Integrated roles in nuclear RNA metabolism and transcription termination. WIREs RNA. 2016 Jan;7(1):91–104.

16. Mitchell P, Petfalski E, Shevchenko A, Mann M, Tollervey D. The Exosome: A Conserved Eukaryotic RNA Processing Complex Containing Multiple 3′→5′ Exoribonucleases. Cell. 1997 Nov 14;91(4):457–66.

17. Allmang C, Kufel J, Chanfreau G, Mitchell P, Petfalski E, Tollervey D. Functions of the exosome in rRNA, snoRNA and snRNA synthesis. EMBO J. 1999 Oct 1;18(19):5399–410.

18. Allmang C, Mitchell P, Petfalski E, Tollervey D. Degradation of ribosomal RNA precursors by the exosome. Nucleic Acids Res. 2000 Apr 15;28(8):1684–91.

19. Malet H, Topf M, Clare DK, Ebert J, Bonneau F, Basquin J, et al. RNA channelling by the eukaryotic exosome. EMBO Rep. 2010 Dec;11(12):936–42.

20. Schilders G, Raijmakers R, Raats JMH, Pruijn GJM. MPP6 is an exosome-associated RNA-binding protein involved in 5.8S rRNA maturation. Nucleic Acids Res. 2005;33(21):6795–804.

21. Mitchell P, Petfalski E, Houalla R, Podtelejnikov A, Mann M, Tollervey D. Rrp47p Is an Exosome-Associated Protein Required for the 3′ Processing of Stable RNAs. Mol Cell Biol. 2003 Oct;23(19):6982–92.

22. Schuch B, Feigenbutz M, Makino DL, Falk S, Basquin C, Mitchell P, et al. The exosome-binding factors Rrp6 and Rrp47 form a composite surface for recruiting the Mtr4 helicase. EMBO J. 2014 Dec;33(23):2829–46.

23. Wasmuth EV, Zinder JC, Zattas D, Das M, Lima CD. Structure and reconstitution of yeast Mpp6-nuclear exosome complexes reveals that mpp6 stimulates RNA decay and recruits the Mtr4 helicase. eLife. 2017;6.

24. LaCava J, Houseley J, Saveanu C, Petfalski E, Thompson E, Jacquier A, et al. RNA degradation by the exosome is promoted by a nuclear polyadenylation complex. Cell. 2005 Jun 3;121(5):713–24.

25. Puno MR, Lima CD. Structural basis for RNA surveillance by the human nuclear exosome targeting (NEXT) complex. Cell. 2022 Jun 9;185(12):2132–2147.e26.

26. Silla T, Schmid M, Dou Y, Garland W, Milek M, Imami K, et al. The human ZC3H3 and RBM26/27 proteins are critical for PAXT-mediated nuclear RNA decay. Nucleic Acids Res. 2020 Mar 18;48(5):2518–30.

27. Meola N, Domanski M, Karadoulama E, Chen Y, Gentil C, Pultz D, et al. Identification of a Nuclear Exosome Decay Pathway for Processed Transcripts. Mol Cell. 2016 Nov 3;64(3):520–33.

28. Chen CY, Gherzi R, Ong SE, Chan EL, Raijmakers R, Pruijn GJM, et al. AU Binding Proteins Recruit the Exosome to Degrade ARE-Containing mRNAs. Cell. 2001 Nov 16;107(4):451–64.

29. Otsuka H, Fukao A, Funakami Y, Duncan KE, Fujiwara T. Emerging Evidence of Translational Control by AU-Rich Element-Binding Proteins. Front Genet [Internet]. 2019 May 2 [cited 2024 Sep 30];10. Available from: https://www.frontiersin.org/journals/genetics/articles/10.3389/fgene.2019.00332/full

30. Puno MR, Weick EM, Das M, Lima CD. SnapShot: The RNA Exosome. Cell. 2019 Sep 19;179(1):282–282.e1.

31. de Amorim J, Slavotinek A, Fasken MB, Corbett AH, Morton DJ. Modeling Pathogenic Variants in the RNA Exosome. RNA Dis Houst Tex. 2020;7:e1166.

32. Lubas M, Christensen MS, Kristiansen MS, Domanski M, Falkenby LG, Lykke-Andersen S, et al. Interaction Profiling Identifies the Human Nuclear Exosome Targeting Complex. Mol Cell. 2011 Aug;43(4):624–37.

33. Wu G, Schmid M, Rib L, Polak P, Meola N, Sandelin A, et al. A Two-Layered Targeting Mechanism Underlies Nuclear RNA Sorting by the Human Exosome. Cell Rep. 2020 Feb 18;30(7):2387–2401.e5.

34. Lubas M, Andersen PR, Schein A, Dziembowski A, Kudla G, Jensen TH. The human nuclear exosome targeting complex is loaded onto newly synthesized RNA to direct early ribonucleolysis. Cell Rep. 2015 Jan 13;10(2):178–92.

35. Gockert M, Schmid M, Jakobsen L, Jens M, Andersen JS, Jensen TH. Rapid factor depletion highlights intricacies of nucleoplasmic RNA degradation. Nucleic Acids Res. 2022 Feb 22;50(3):1583–600.

36. EXOSC3 DepMap Gene Summary [Internet]. [cited 2024 May 29]. Available from: https://depmap.org/portal/gene/EXOSC3?tab=overview

37. Fasken MB, Losh JS, Leung SW, Brutus S, Avin B, Vaught JC, et al. Insight into the RNA Exosome Complex Through Modeling Pontocerebellar Hypoplasia Type 1b Disease Mutations in Yeast. Genetics. 2017 Jan 1;205(1):221–37.

38. Morton DJ, Jalloh B, Kim L, Kremsky I, Nair RJ, Nguyen KB, et al. A Drosophila model of Pontocerebellar Hypoplasia reveals a critical role for the RNA exosome in neurons. PLoS Genet. 2020 Jul;16(7):e1008901.

39. Sterrett MC, Cureton LA, Cohen LN, van Hoof A, Khoshnevis S, Fasken MB, et al. Comparative analyses of disease-linked missense mutations in the RNA exosome modeled in budding yeast reveal distinct functional consequences in translation. bioRxiv. 2023 Oct 19;2023.10.18.562946.

40. Gillespie A, Gabunilas J, Jen JC, Chanfreau GF. Mutations of EXOSC3/Rrp40p associated with neurological diseases impact ribosomal RNA processing functions of the exosome in S. cerevisiae. RNA. 2017 Apr;23(4):466–72.

41. Falk S, Bonneau F, Ebert J, Kögel A, Conti E. Mpp6 Incorporation in the Nuclear Exosome Contributes to RNA Channeling through the Mtr4 Helicase. Cell Rep. 2017 Sep 5;20(10):2279–86.

42. Müller JS, Burns DT, Griffin H, Wells GR, Zendah RA, Munro B, et al. RNA exosome mutations in pontocerebellar hypoplasia alter ribosome biogenesis and p53 levels. Life Sci Alliance. 2020 Aug;3(8):e202000678.

43. Weick EM, Puno MR, Januszyk K, Zinder JC, DiMattia MA, Lima CD. Helicase-Dependent RNA Decay Illuminated by a Cryo-EM Structure of a Human Nuclear RNA Exosome-MTR4 Complex. Cell. 2018 Jun 14;173(7):1663–1677.e21.

44. Latini C, Eichlinger J, Fuchs AL, Zhai SN, Ho-Xuan H, Lehmann G, et al. Cytoplasmic DIS3 is an exosome-independent endoribonuclease with catalytic activity toward circular RNAs. Cell Rep. 2025 Jun;44(6):115769.

45. McCracken NA, Justice SAP, Smith-Kinnaman WR, Wijeratne HS, Runnebohm AM, Pelletier S, et al. Application of Whole Proteome Thermal Shift Assays to Define PERK-dependent Changes in Protein Homeostasis during the Unfolded Protein Response [Internet]. bioRxiv; 2025 [cited 2025 May 7]. p. 2025.04.08.647882. Available from: https://www.biorxiv.org/content/10.1101/2025.04.08.647882v1

46. Gaetani M, Sabatier P, Saei AA, Beusch CM, Yang Z, Lundström SL, et al. Proteome Integral Solubility Alteration: A High-Throughput Proteomics Assay for Target Deconvolution. J Proteome Res. 2019 Nov 1;18(11):4027–37.

47. Gaetani M, Zubarev RA. Proteome Integral Solubility Alteration (PISA) for High-Throughput Ligand Target Deconvolution with Increased Statistical Significance and Reduced Sample Amount. Methods Mol Biol Clifton NJ. 2023;2554:91–106.

48. Berlivet S, Moussette S, Ouimet M, Verlaan DJ, Koka V, Al Tuwaijri A, et al. Interaction between genetic and epigenetic variation defines gene expression patterns at the asthma-associated locus 17q12-q21 in lymphoblastoid cell lines. Hum Genet. 2012 Jul;131(7):1161–71.

49. Gicquel C, Rossignol S, Cabrol S, Houang M, Steunou V, Barbu V, et al. Epimutation of the telomeric imprinting center region on chromosome 11p15 in Silver-Russell syndrome. Nat Genet. 2005 Sep;37(9):1003–7.

50. Jentarra GM, Rice SG, Olfers S, Rajan C, Saffen DM, Narayanan V. Skewed allele-specific expression of the NF1 gene in normal subjects: a possible mechanism for phenotypic variability in neurofibromatosis type 1. J Child Neurol. 2012 Jun;27(6):695–702.

51. St. Pierre CL, Macias-Velasco JF, Wayhart JP, Yin L, Semenkovich CF, Lawson HA. Genetic, epigenetic, and environmental mechanisms govern allele-specific gene expression. Genome Res. 2022 Jun;32(6):1042–57.

52. Pastinen T. Genome-wide allele-specific analysis: insights into regulatory variation. Nat Rev Genet. 2010 Aug;11(8):533–8.

53. Hinnebusch AG, Ivanov IP, Sonenberg N. Translational control by 5’-untranslated regions of eukaryotic mRNAs. Science. 2016 Jun 17;352(6292):1413–6.

54. Leppek K, Das R, Barna M. Functional 5′ UTR mRNA structures in eukaryotic translation regulation and how to find them. Nat Rev Mol Cell Biol. 2018 Mar;19(3):158–74.

55. Mayr C. Regulation by 3’-Untranslated Regions. Annu Rev Genet. 2017 Nov 27;51:171–94.

56. Dassi E, Re A, Leo S, Tebaldi T, Pasini L, Peroni D, et al. AURA 2: Empowering discovery of post-transcriptional networks. Transl Austin Tex. 2014;2(1):e27738.

57. Dassi E, Malossini A, Re A, Mazza T, Tebaldi T, Caputi L, et al. AURA: Atlas of UTR Regulatory Activity. Bioinformatics. 2012 Jan 1;28(1):142–4.

58. Juszkiewicz S, Hegde RS. Quality Control of Orphaned Proteins. Mol Cell. 2018 Aug 2;71(3):443–57.

59. Peck Justice SA, Barron MP, Qi GD, Wijeratne HRS, Victorino JF, Simpson ER, et al. Mutant thermal proteome profiling for characterization of missense protein variants and their associated phenotypes within the proteome. J Biol Chem. 2020 Nov;295(48):16219–38.

60. Mateus A, Kurzawa N, Becher I, Sridharan S, Helm D, Stein F, et al. Thermal proteome profiling for interrogating protein interactions. Mol Syst Biol [Internet]. 2020 Mar [cited 2022 Feb 18];16(3). Available from: https://onlinelibrary.wiley.com/doi/10.15252/msb.20199232

61. Singh S, Vanden Broeck A, Miller L, Chaker-Margot M, Klinge S. Nucleolar maturation of the human small subunit processome. Science. 2021 Sep 10;373(6560):eabj5338.

62. Phipps KR, Charette JM, Baserga SJ. The SSU Processome in Ribosome Biogenesis – Progress and Prospects. Wiley Interdiscip Rev RNA. 2011 Jan;2(1):1– 21.

63. Costello JL, Stead JA, Feigenbutz M, Jones RM, Mitchell P. The C-terminal region of the exosome-associated protein Rrp47 is specifically required for box C/D small nucleolar RNA 3’-maturation. J Biol Chem. 2011 Feb 11;286(6):4535–43.

64. Milligan L, Decourty L, Saveanu C, Rappsilber J, Ceulemans H, Jacquier A, et al. A Yeast Exosome Cofactor, Mpp6, Functions in RNA Surveillance and in the Degradation of Noncoding RNA Transcripts. Mol Cell Biol. 2008 Sep;28(17):5446–57.

65. Halevy A, Lerer I, Cohen R, Kornreich L, Shuper A, Gamliel M, et al. Novel EXOSC3 mutation causes complicated hereditary spastic paraplegia. J Neurol. 2014 Nov;261(11):2165–9.

66. Ryan MM, Cooke-Yarborough CM, Procopis PG, Ouvrier RA. Anterior horn cell disease and olivopontocerebellar hypoplasia. Pediatr Neurol. 2000 Aug;23(2):180–4.

67. Mu W, Heller T, Barañano KW. Two siblings with a novel variant of EXOSC3 extended phenotypic spectrum of pontocerebellar hypoplasia 1B to an exceptionally mild form. BMJ Case Rep CP. 2021 Jan 1;14(1):e236732.

68. Kuliyev E, Gingras S, Guy CS, Howell S, Vogel P, Pelletier S. Overlapping Role of SCYL1 and SCYL3 in Maintaining Motor Neuron Viability. J Neurosci Off J Soc Neurosci. 2018 Mar 7;38(10):2615–30.

69. Pelletier S, Gingras S, Green DR. Mouse genome engineering via CRISPR-Cas9 for study of immune function. Immunity. 2015 Jan 20;42(1):18–27.

70. SourceForge [Internet]. 2023 [cited 2024 Jun 12]. BBMap. Available from: https://sourceforge.net/projects/bbmap/

71. Babraham Bioinformatics - FastQC A Quality Control tool for High Throughput Sequence Data [Internet]. [cited 2024 Jun 12]. Available from: https://www.bioinformatics.babraham.ac.uk/projects/fastqc/

72. Yates AD, Achuthan P, Akanni W, Allen J, Allen J, Alvarez-Jarreta J, et al. Ensembl 2020. Nucleic Acids Res. 2020 Jan 8;48(D1):D682–8.

73. Dobin A, Davis CA, Schlesinger F, Drenkow J, Zaleski C, Jha S, et al. STAR: ultrafast universal RNA-seq aligner. Bioinforma Oxf Engl. 2013 Jan 1;29(1):15–21.

74. Liao Y, Smyth GK, Shi W. featureCounts: an efficient general purpose program for assigning sequence reads to genomic features. Bioinforma Oxf Engl. 2014 Apr 1;30(7):923–30.

75. Robinson MD, McCarthy DJ, Smyth GK. edgeR: a Bioconductor package for differential expression analysis of digital gene expression data. Bioinforma Oxf Engl. 2010 Jan 1;26(1):139–40.

76. Stephens M. False discovery rates: a new deal. Biostat Oxf Engl. 2017 Apr 1;18(2):275–94.

77. Ge SX, Jung D, Yao R. ShinyGO: a graphical gene-set enrichment tool for animals and plants. Bioinformatics. 2020 Apr 15;36(8):2628–9.

78. Snel B, Lehmann G, Bork P, Huynen MA. STRING: a web-server to retrieve and display the repeatedly occurring neighbourhood of a gene. Nucleic Acids Res. 2000 Sep 15;28(18):3442–4.

79. von Mering C, Huynen M, Jaeggi D, Schmidt S, Bork P, Snel B. STRING: a database of predicted functional associations between proteins. Nucleic Acids Res. 2003 Jan 1;31(1):258–61.

80. von Mering C, Jensen LJ, Snel B, Hooper SD, Krupp M, Foglierini M, et al. STRING: known and predicted protein-protein associations, integrated and transferred across organisms. Nucleic Acids Res. 2005 Jan 1;33(Database issue):D433-437.

81. Szklarczyk D, Gable AL, Lyon D, Junge A, Wyder S, Huerta-Cepas J, et al. STRING v11: protein-protein association networks with increased coverage, supporting functional discovery in genome-wide experimental datasets. Nucleic Acids Res. 2019 Jan 8;47(D1):D607–13.

82. Szklarczyk D, Kirsch R, Koutrouli M, Nastou K, Mehryary F, Hachilif R, et al. The STRING database in 2023: protein-protein association networks and functional enrichment analyses for any sequenced genome of interest. Nucleic Acids Res. 2023 Jan 6;51(D1):D638–46.

83. Wickham H. ggplot2: Elegant Graphics for Data Analysis [Internet]. New York, NY: Springer; 2009 [cited 2023 Jun 6]. Available from: https://link.springer.com/10.1007/978-0-387-98141-3

84. Zinder JC, Lima CD. Targeting RNA for processing or destruction by the eukaryotic RNA exosome and its cofactors. Genes Dev. 2017 Jan 15;31(2):88–100.

85. Venny 2.1.0 [Internet]. [cited 2025 May 27]. Available from: https://bioinfogp.cnb.csic.es/tools/venny/

86. Metsalu T, Vilo J. ClustVis: a web tool for visualizing clustering of multivariate data using Principal Component Analysis and heatmap. Nucleic Acids Res. 2015 Jul 1;43(W1):W566–70.

