## Supplementary material for "EXOSC3 S1-domain variants implicated in PCH1b alter RNA exosome cap subunit abundance and thermal stability disrupting rRNA processing and targeting of AU-rich mRNA": Figure S1

A

| Variants in Humans | Pathogenicity | Functional Domain | Allele Frequency (gnomAD) | Pathogenic Variant Models |
|---|---|---|---|---|
| G31A | Pathogenic | N-terminal Domain | 1.12e-5 | 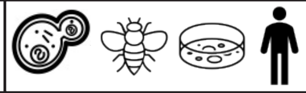 |
| D132A | Pathogenic | S1 Domain | 7.06e-4 | 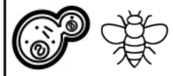 |
| G135R | VUS | S1 Domain | 5.58e-6 | |
| A139P | Pathogenic | S1 Domain | 6.20e-6 | |
| G191D | Likely Pathogenic | interdomain | 1.43e-5 | |
| G191C | Pathogenic | interdomain | not available | 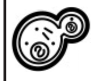 |
| W238R | Pathogenic | KH Domain | not available | 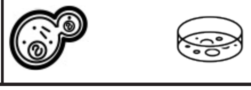 |

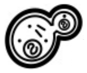 yeast 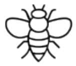 fly 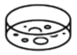 cultured murine cells 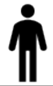 patient derived cells

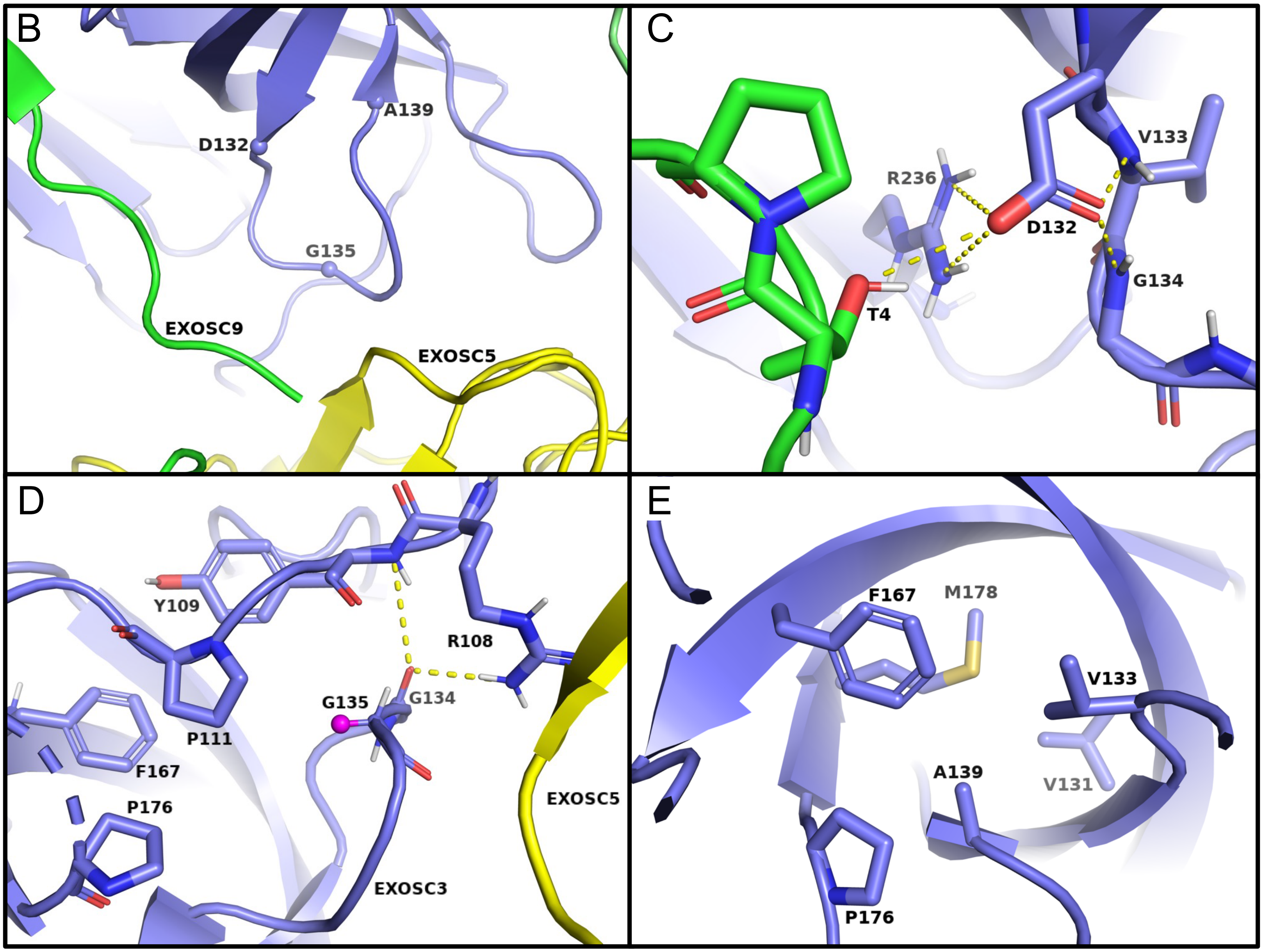
