## Supplementary figures and images for "EXOSC3 S1-domain variants implicated in PCH1b alter RNA exosome cap subunit abundance and thermal stability disrupting rRNA processing and targeting of AU-rich mRNA"

### Figure S2

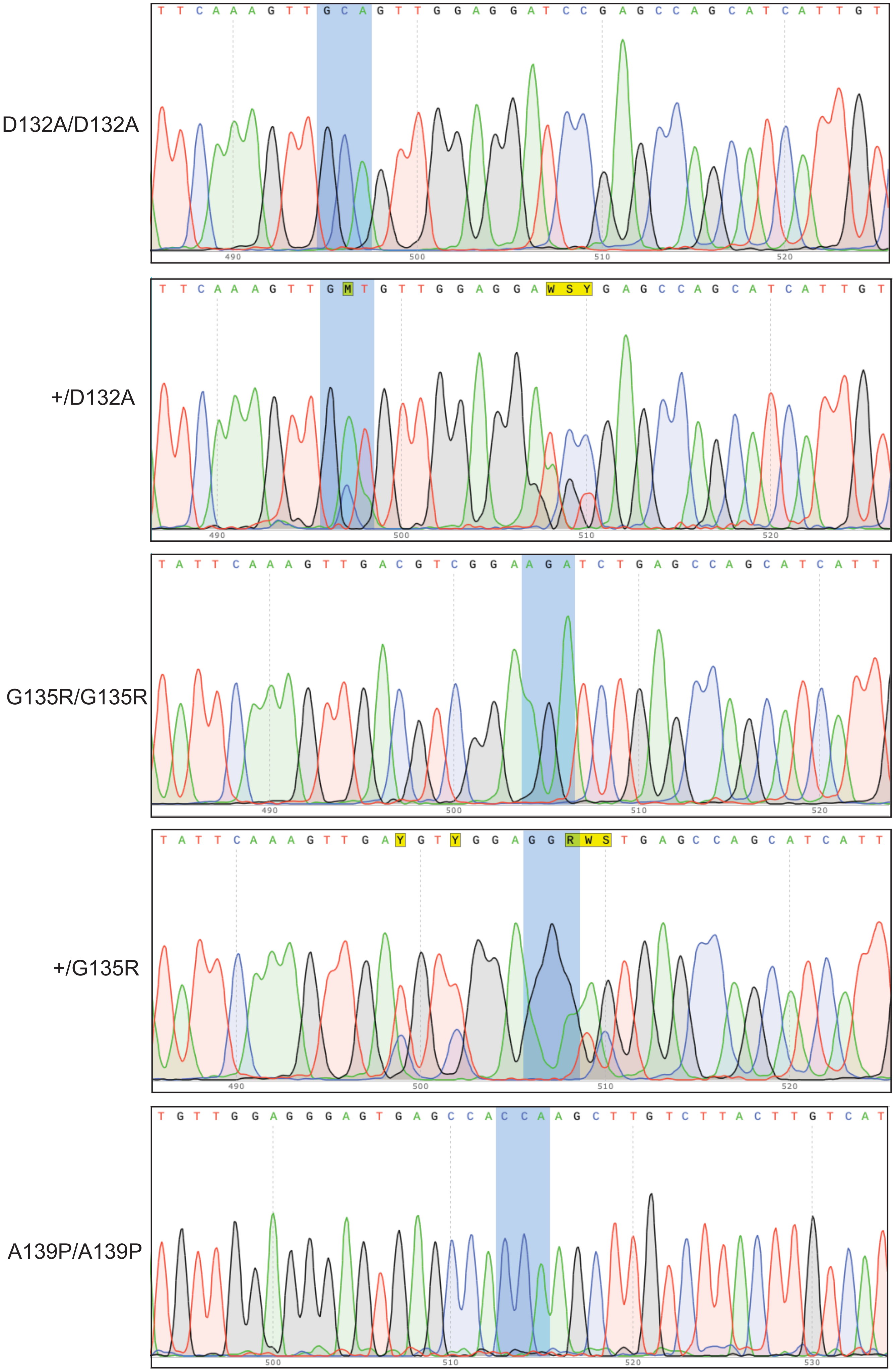

### Figure S3

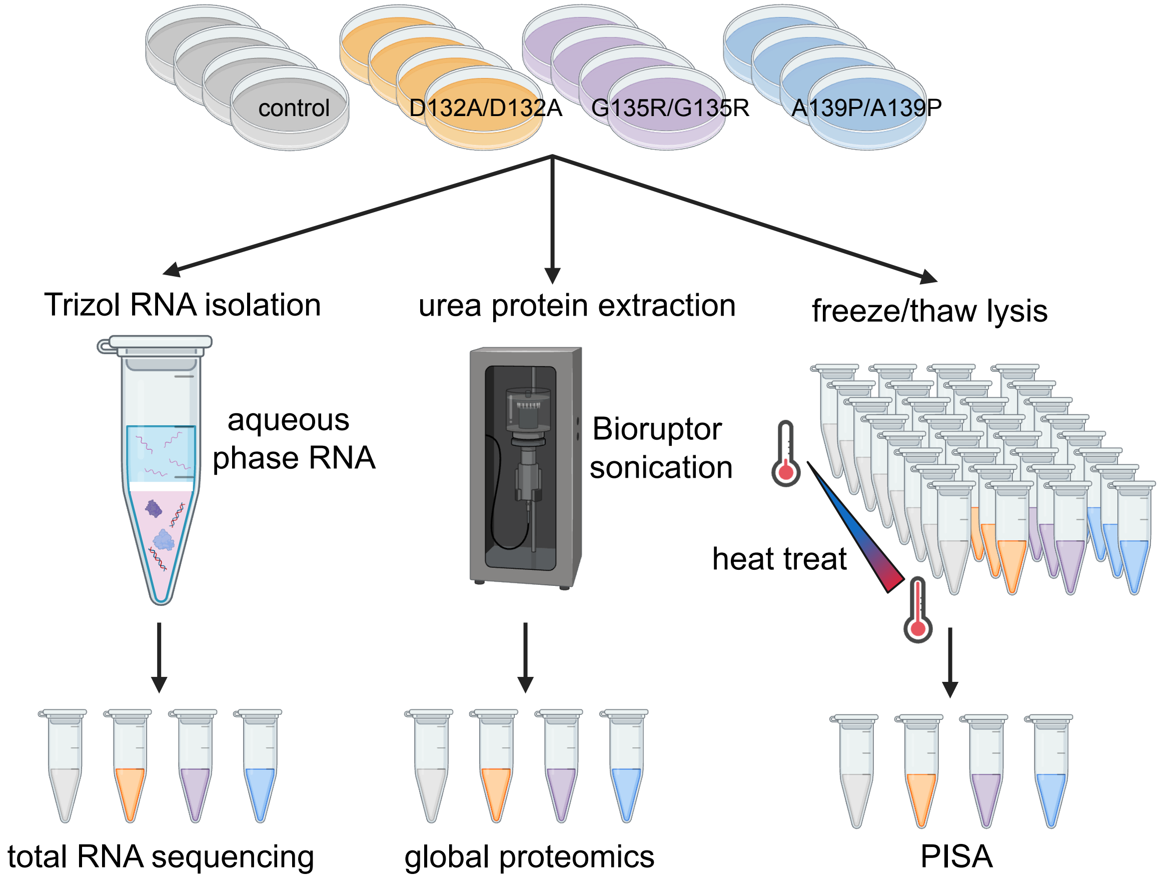

### Figure S4

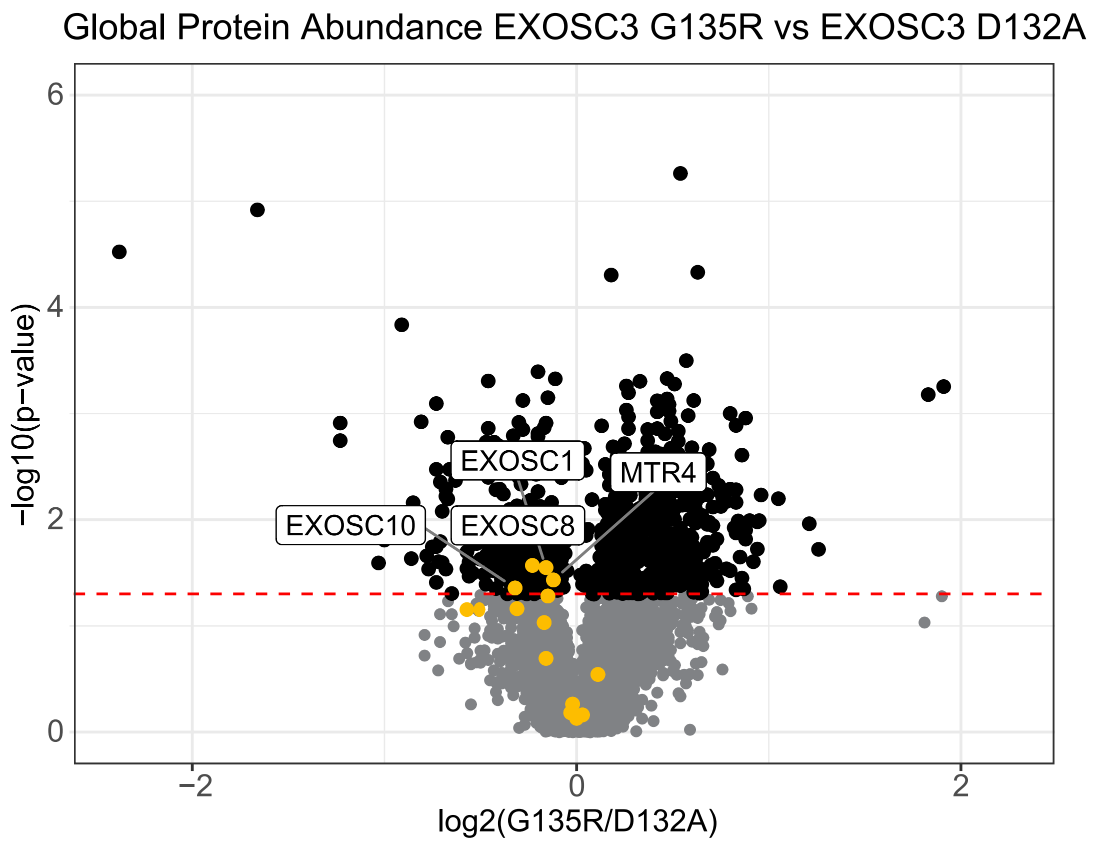

### Figure S5

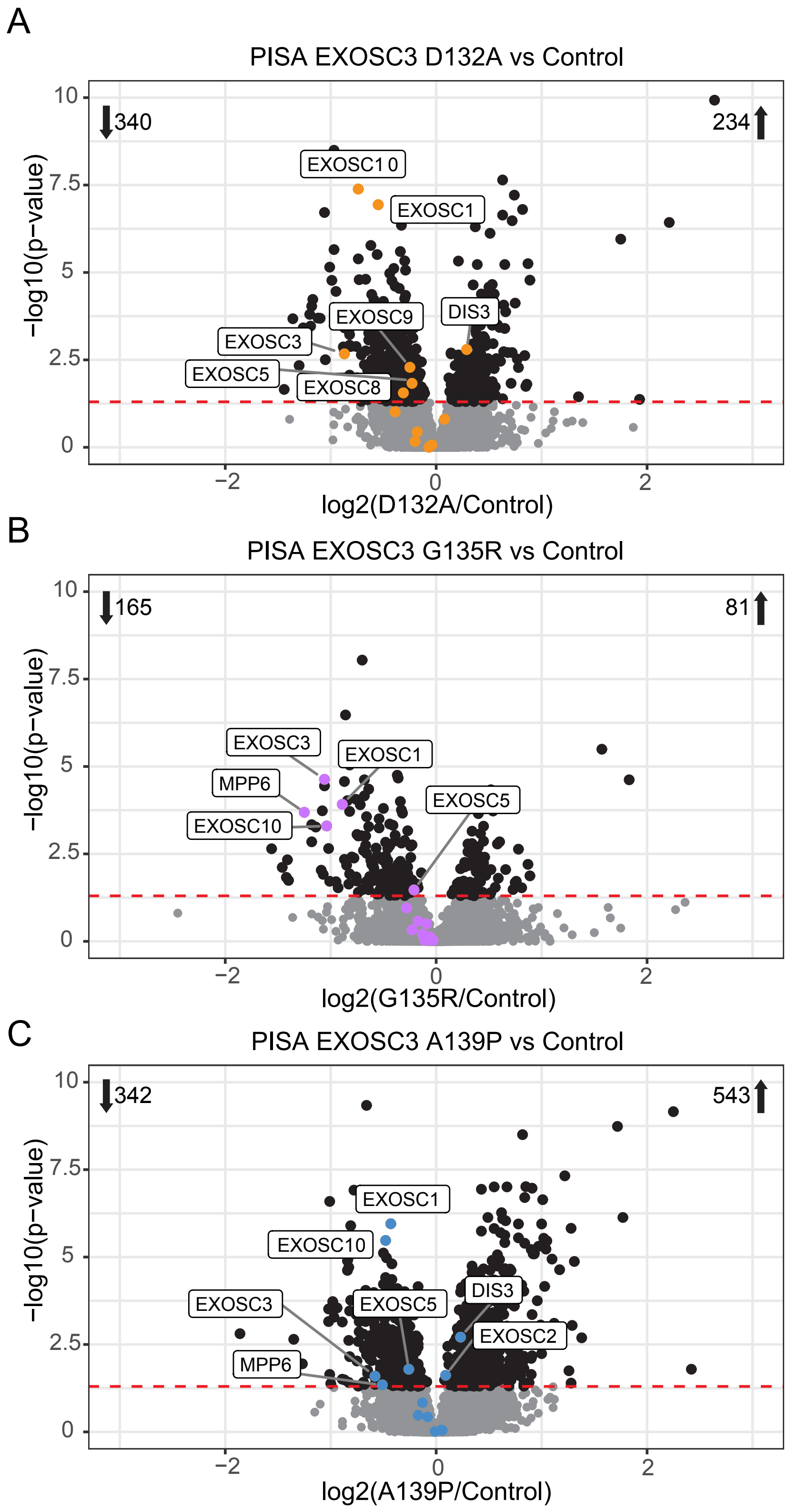

### Figure S6

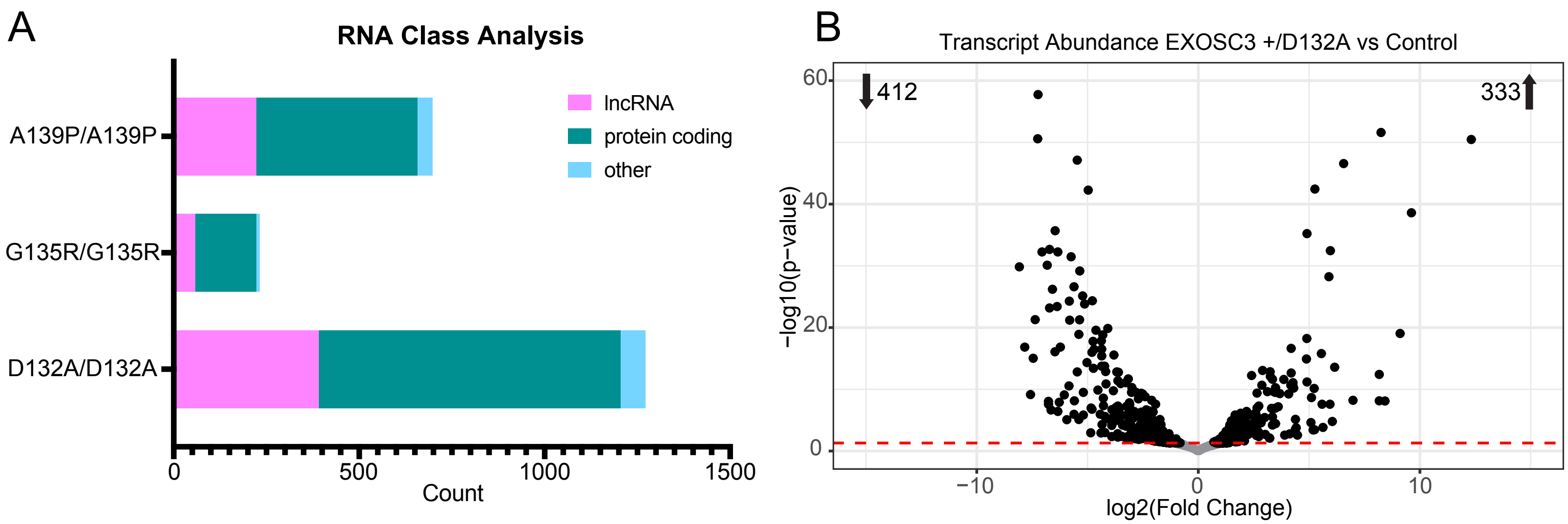

### Figure S7

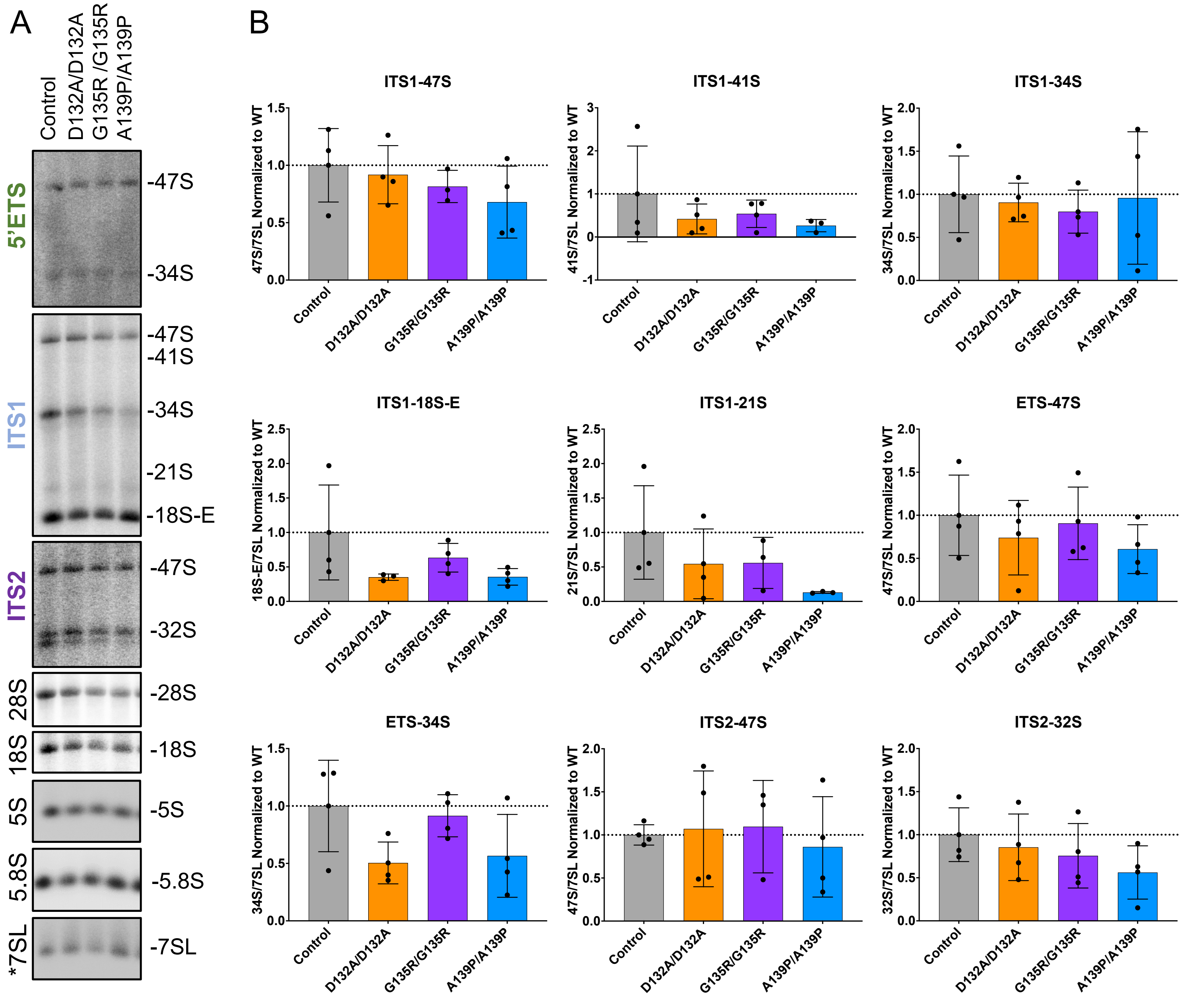

### Figure S8

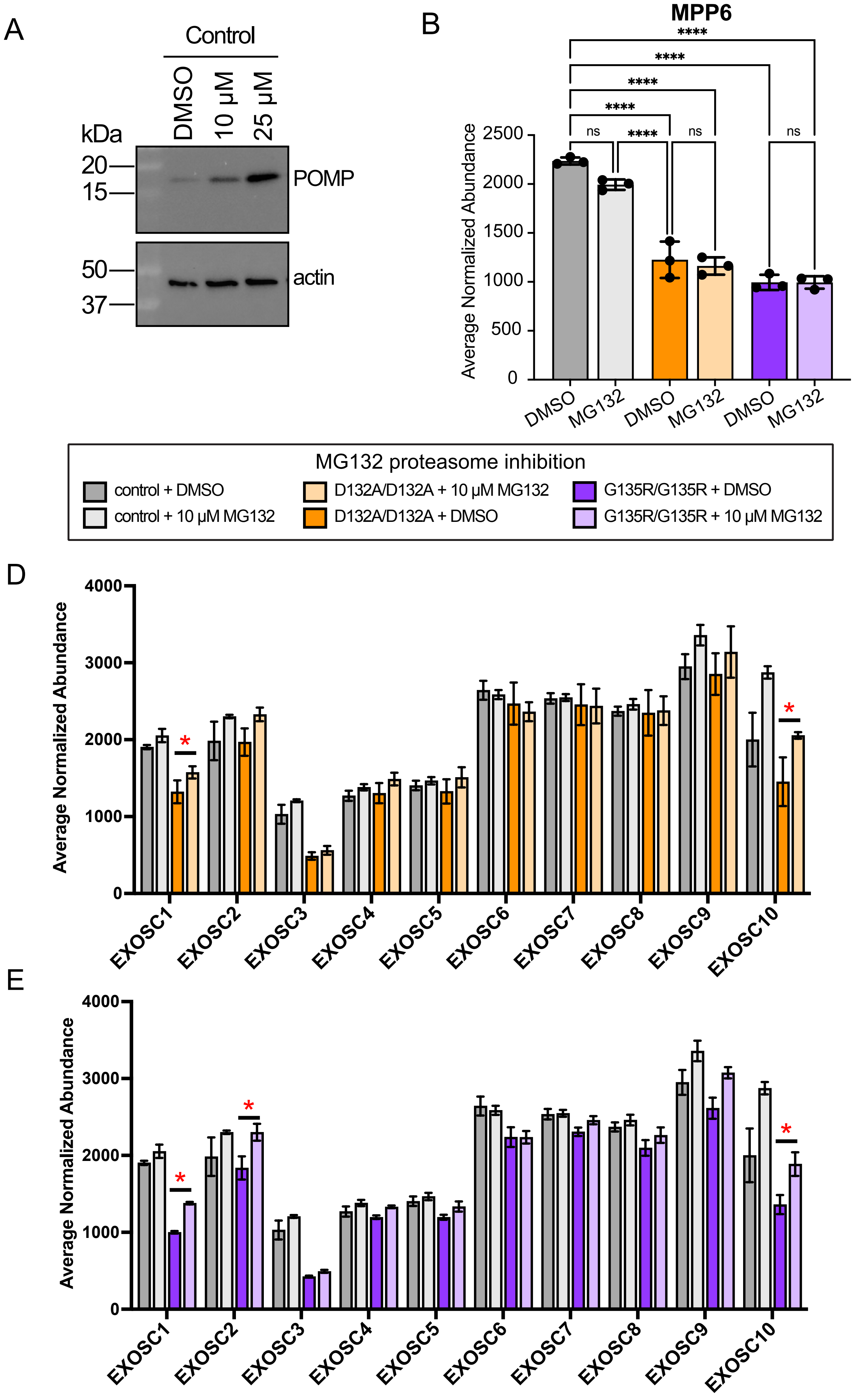
